# MicrobiomePhylo: A New Tool for Metabarcoding Data Downstream Analysis - A Real-World Data Analysis Demonstration

**DOI:** 10.1101/2024.04.08.588598

**Authors:** Camilla Veronica Tafuro

**Affiliations:** Independent researcher

## Abstract

The field of microbiome research has rapidly expanded, driven by advancements in sequencing technologies that generate vast amounts of data. To navigate this complexity, MicrobiomePhylo has been developed as a web platform tailored for the downstream analysis of microbiome amplicon sequencing data. It stands out for its ability to process QIIME2 artifact files (.qza), offering an intuitive interface, extensive analytical functions, and advanced visualization features, making it indispensable for bioinformaticians and biologists alike. MicrobiomePhylo enhances the understanding of complex datasets by facilitating detailed analysis and visualization. Its effectiveness is showcased through the analysis of the “Moving Pictures” tutorial dataset from QIIME2, a benchmark in microbiome research, featuring 16S rRNA gene sequencing data from various sample types over time.

The platform supports a comprehensive workflow including data upload, pre-processing, refinement, visualization, exploration, and advanced statistical analysis. Features include data filtering, rarefaction, and diverse visualizations like barplots, boxplots, and heatmaps. Advanced analyses cover alpha and beta diversity, abundance analysis, differential abundance analysis, and correlation studies, backed by robust statistical methods.

A demonstration with the “Moving Pictures” dataset highlights MicrobiomePhylo’s capability to manage real-world data. Customization options for variables, taxonomic levels, plot dimensions, and group highlighting improve result interpretability and visual appeal. Analysis at the phylum level across different body sites reveals insights into microbial composition and diversity. The generated visualizations distinguish microbial community compositions and taxa abundance between body sites, emphasizing the site-specific nature of the human microbiome.

**Importance:** The significance of MicrobiomePhylo in microbiome research is profound. As sequencing technologies advance, generating vast datasets, researchers face the challenge of analyzing this data efficiently. MicrobiomePhylo, a web platform designed for microbiome amplicon sequencing data analysis, directly processes QIIME2 artifact files (.qza), offering an accessible interface for both bioinformaticians and biologists. Its comprehensive analytical and advanced visualization capabilities are crucial for understanding complex datasets, enabling deeper insights into microbial composition and diversity. The platform supports a full workflow from data upload to advanced statistical analysis, including data filtering, rarefaction, and various visualizations for taxa abundances, correlations with continuous variables, diversity and more. This facilitates a thorough investigation of microbiome data, making MicrobiomePhylo an indispensable tool in the field, enhancing our understanding of the microbiome through specialized, user-friendly analysis capabilities.

## Introduction

The field of microbiome research is advancing at an unprecedented pace, driven by the critical need for precise analysis and visualization of complex datasets. <u>MicrobiomePhylo</u>, a state-of-the-art web platform, addresses this need by providing an extensive array of tools for the downstream analysis of microbiome amplicon sequencing data [1]. Engineered to seamlessly process QIIME2 [2] artifact files (.qza), the platform distinguishes itself through its intuitive interface, comprehensive analytical functions, and advanced visualization features, making it an indispensable tool for both bioinformaticians and biologists.

Microbiome research plays a pivotal role in unraveling the complex interactions between microbial communities and their hosts, shedding light on their significant implications for health and disease, thereby underscoring the critical need for advanced platforms like MicrobiomePhylo to facilitate in-depth analysis and visualization of microbiome data [4]. To showcase the platform’s prowess, we present a detailed analysis using the “Moving Pictures” tutorial dataset from the QIIME2 analysis package, a benchmark dataset in the microbiome research community.

The “Moving Pictures” dataset provides an ideal testbed for demonstrating MicrobiomePhylo’s capabilities [1]. It comprises 16S rRNA gene sequencing data derived from four sample types (two human hosts) collected at multiple time points, offering a dynamic view of the microbiome’s temporal and spatial variability. This dataset allows researchers to explore a wide range of analytical questions, from basic compositional analysis to complex comparative and temporal studies.

### Analytical Workflow with MicrobiomePhylo: A Comprehensive Guide

#### Data Upload and Preprocessing

The initial step involves uploading QIIME2 [2] artifacts and metadata files. MicrobiomePhylo ensures a streamlined analysis onset by enforcing format compliance and specific naming conventions, thereby optimizing compatibility and user experience.

#### Data Refinement and Standardization

MicrobiomePhylo provides tools [3] for adjusting detection and prevalence thresholds and excluding specific samples from analyses, allowing users to refine their dataset based on specific research needs. This step is vital for focusing the analysis on relevant data points and minimizing the impact of outliers or noise. The option for rarefaction standardizes sampling effort across samples, a critical step for comparative analyses. This is particularly important in comparative analyses, as it ensures that differences observed between samples are not merely artifacts of varying sequencing efforts. Together, data filtering and rarefaction are critical for preparing the dataset in a way that facilitates meaningful and fair comparisons across samples.

#### Visualization and Exploration

The platform facilitates an immersive exploration of data. It enables users to visualize the reads distribution and explore the microbial composition through various plots, including barplots, boxplots, and heatmaps. These visualizations are customizable, allowing for tailored analyses based on grouping variables and other parameters. By leveraging these tools, researchers can gain an intuitive understanding of microbial community structures and distributions, fostering hypothesis generation and the identification of significant patterns warranting further investigation.

#### Advanced Statistical Analysis

MicrobiomePhylo goes beyond basic visualization, offering advanced analyses such as alpha and beta diversity assessments, abundance analysis, differential abundance analysis based on LEfSe and correlation studies. These analyses are supported by robust statistical methodologies, including the Wilcoxon test and PERMANOVA [5][6], providing users with the means to uncover significant patterns and relationships within their data. This step is crucial for identifying statistically significant differences and associations within the microbiome data, ultimately leading to insights that can drive scientific discovery and inform future research directions.

#### Demonstration with the “Moving Pictures” Dataset

Utilizing the “Moving Pictures” dataset, MicrobiomePhylo showcases its adeptness at managing real-world microbiome data with with precision and ease. By bridging the gap between complex data analysis and intuitive, accessible tools, MicrobiomePhylo is poised to advance the field of microbiome research, enabling discoveries that deepen our understanding of the microbial world.

#### Customization Features

MicrobiomePhylo provides the tools to visualize your data in ways that best suit your research objectives with Customization Options to enhance the interpretability and aesthetic appeal of research results. These includes:

#### Grouping Variables

Tailor the analysis by selecting grouping variables that align with the research questions. This feature enables users to compare microbial communities based on factors such as treatment, location, or any other relevant categorization.

#### Taxonomic Levels

Choose the taxonomic resolution that best fits the analysis, from phylum down to species level. This flexibility allows for both broad overviews and detailed investigations of microbial composition.

#### Plot Dimensions

Adjust the dimensions of each plot to optimize visualization for presentations, publications, or in-depth analysis, catering to diverse analytical needs.

#### Group Highlighting

Assign specific colors to different groups or conditions within the dataset, making it easier to distinguish and compare these groups across various visualizations. Once a color palette is selected, it is consistently applied across all relevant visualizations within your analysis, ensuring a cohesive and professional appearance of your output. Color customization not only improves the visual appeal of the analyses but also enhances interpretability by providing clear visual cues that help distinguish between different microbial communities or experimental conditions.

## Methods

In this study, we utilized the MicrobiomePhylo web application to conduct a comprehensive analysis of microbiome data derived from the Moving Pictures tutorial dataset. This dataset comprises a feature table, taxonomy table, and a rooted phylogenetic tree, all in QIIME2 artifact format (.qza), accompanied by metadata in TSV format. Our analysis adhered to a structured methodology, concentrating on the phylum taxonomic level and facilitating comparisons across different body sites.

### Data Preparation and Filtering

Initially, the dataset underwent a filtering process to remove features detected in less than 5% of the samples and with a detection threshold established at 1.

No samples were excluded from this analysis.

To ensure uniformity in sampling depth across the dataset, rarefaction was performed at the minimum read count per sample, allowing for the inclusion of all samples.

### Analytical Workflow

#### Rarefaction

Data was rarefied to ensure even sampling depth across all samples, thus minimizing bias in diversity estimates [7].

#### Abundance Analysis

The rarefied data was normalized to relative abundance to accurately represent the proportional distribution of taxa within each sample [8][9].

#### Correlation Analysis

We explored the associations between microbial taxa and metadata variables using Spearman rho correlation. This non-parametric method is apt for datasets not adhering to a normal distribution, assessing monotonic relationships to elucidate how taxa abundance correlates with environmental or physiological factors.

#### Significance Testing

Significant Spearman correlations were identified, and p-values were calculated. To account for multiple comparisons, p-values were adjusted using the BenjaminiHochberg procedure for False Discovery Rate (FDR) control [10].

#### Alpha Diversity Analysis

Within-sample diversity was estimated using metrics such as Observed Species, Chao1, Simpson, and Shannon indices [11].

#### Beta Diversity Analysis

Diversity between samples was assessed using metrics (Jaccard, Bray-Curtis, Unifrac, and Weighted Unifrac) [6] that reflect compositional and phylogenetic differences. PERMANOVA and Betadisper tests were utilized to examine homogeneity of variance [12].

#### Differential Abundance Analysis

LEfSe (Linear discriminant analysis Effect Size) was employed to identify biomarkers and statistically significant differences in taxa abundance across groups [13].

### Statistical Tests and Effect Size Analysis

Wilcoxon test was performed to assess the significance of differences in abundance between groups for the top four taxa. Kruskal-Wallis tests were performed to identify differences in alpha diversity metrics across groups, followed by Dunn tests for post-hoc analysis [14]. Effect sizes were calculated to quantify the magnitude of observed differences.

ANOVA tests for dispersions and permutation tests for pairwise comparisons, followed by Tukey test, were conducted to further explore beta diversity.

### Visualizations

A range of visualizations were generated to visually explore and communicate the complex patterns within the microbiome data. Visual tools are instrumental in revealing insights that might not be immediately apparent from statistical tests alone [15].

#### Reads Distribution

Visualized both overall and per-group to assess sequencing depth.

#### Alpha and Beta Diversity

Rarefaction curves, NMDS plots for various distance metrics, and PCoA plots with scree plots were generated to visualize diversity patterns.

#### Abundance Analysis

Relative abundance barplots, boxplots for top taxa, and detailed heatmaps were created to explore taxa distribution.

#### Significance Annotations

Alpha diversity boxplots were annotated with significance levels to underscore differences between groups.

#### LEfSe Results

Visualized through barplots, abundance plots, and heatmaps to identify and display biomarkers across groups.

This methodical approach facilitated a thorough exploration of the microbiome dataset at the phylum level, enabling insightful comparisons based on body site. The combination of advanced statistical analyses and diverse visualizations provided a comprehensive understanding of the microbial communities present, their diversity, and their potential associations with different body sites.

## Results

Our comprehensive analysis of 34 microbiome samples has yielded insightful findings on the microbial composition and diversity across different body sites. The data summary highlights a range of sequencing reads per sample, from a minimum of 892 to a maximum of 9231, with a total of 149172 reads. The average number of reads was 4387.4, with a median of 3743.5. Detailed below are the results from our various analyses, including diversity assessments, statistical tests, and visualizations of microbial abundance and diversity.

### Reads Distribution per Sample

The distribution of sequencing reads across samples revealed variability in sequencing depth [18], which is crucial for understanding the coverage and reliability of the detected microbial communities.

The variability in sequencing depth highlighted by Fig 1 underscores the importance of normalization methods like rarefaction in microbiome studies. This ensures that comparisons across samples are not biased by differences in sequencing effort. The wide range of sequencing depths also suggests that some samples may have richer microbial diversity captured than others, potentially influencing the observed microbial composition and diversity.

**Fig 1.**
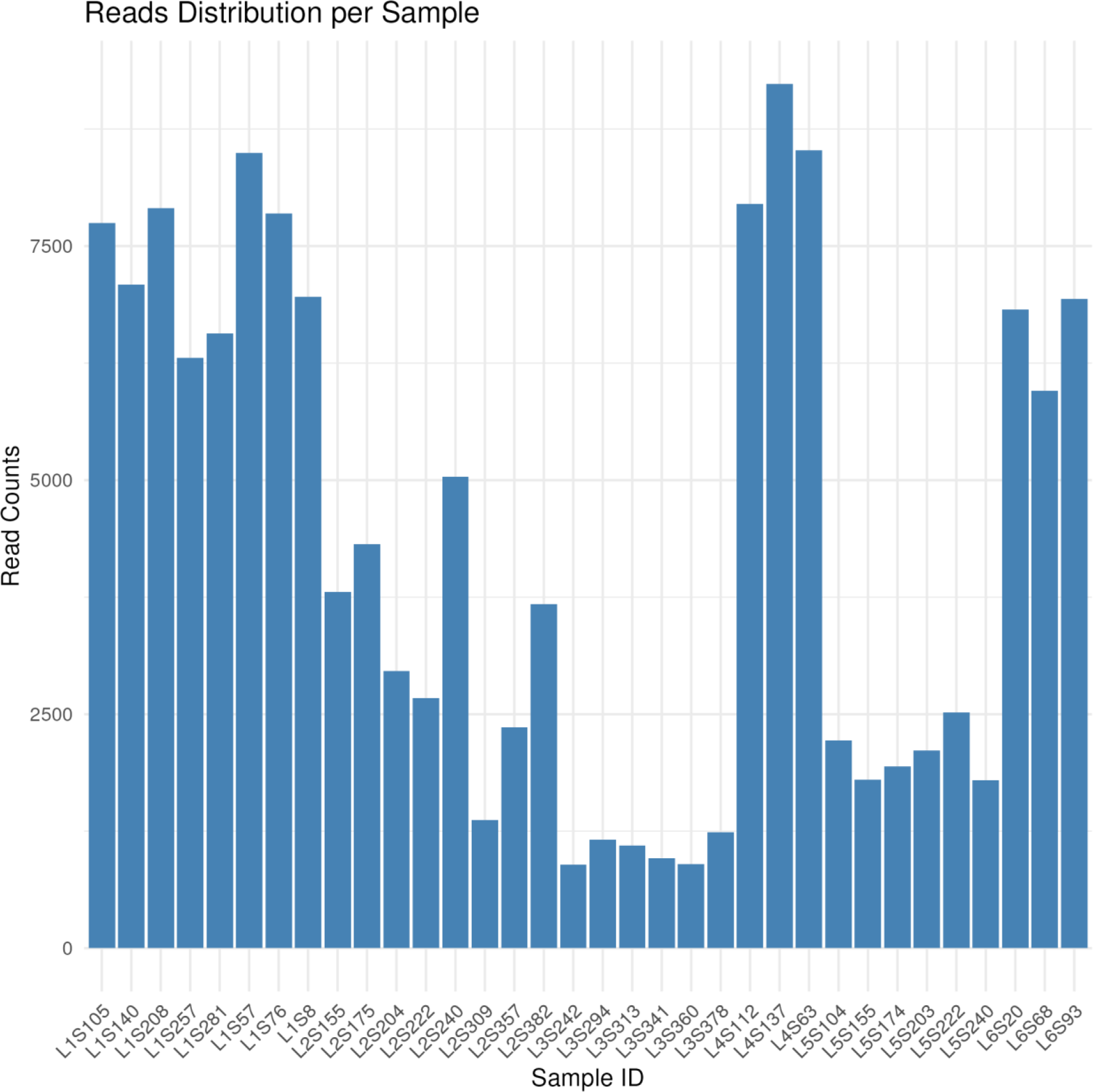
Distribution of Sequencing Reads Across Samples: This plot visualizes the range of sequencing depths achieved across the 34 samples, highlighting the minimum of 892 reads to a maximum of 9231 reads per sample.

It is crucial to determine the point at which additional sequencing does not significantly alter the observed community structure, known as the “plateau” or “saturation” point. Achieving this point is critical for ensuring that the sequencing depth is sufficient to capture the majority of the microbial diversity present, thereby enhancing the reliability of the findings.

Rarefaction was applied at the minimum number of reads per sample at which all observed diversity curves reached a plateau. This plateau indicates that additional sequencing would likely not reveal more diversity within the samples, suggesting that the chosen rarefaction depth captures the majority of the microbial diversity present. Applying rarefaction at this level ensures that comparisons across samples are made on an equal footing, mitigating the potential bias introduced by differences in sequencing effort.

### Abundance Barplot per Group

Our analysis generated abundance barplots for each group, illustrating the relative abundance of microbial taxa across different body sites. These visualizations provide a clear overview of the dominant microbial communities in each sample group.

**Fig 2.**
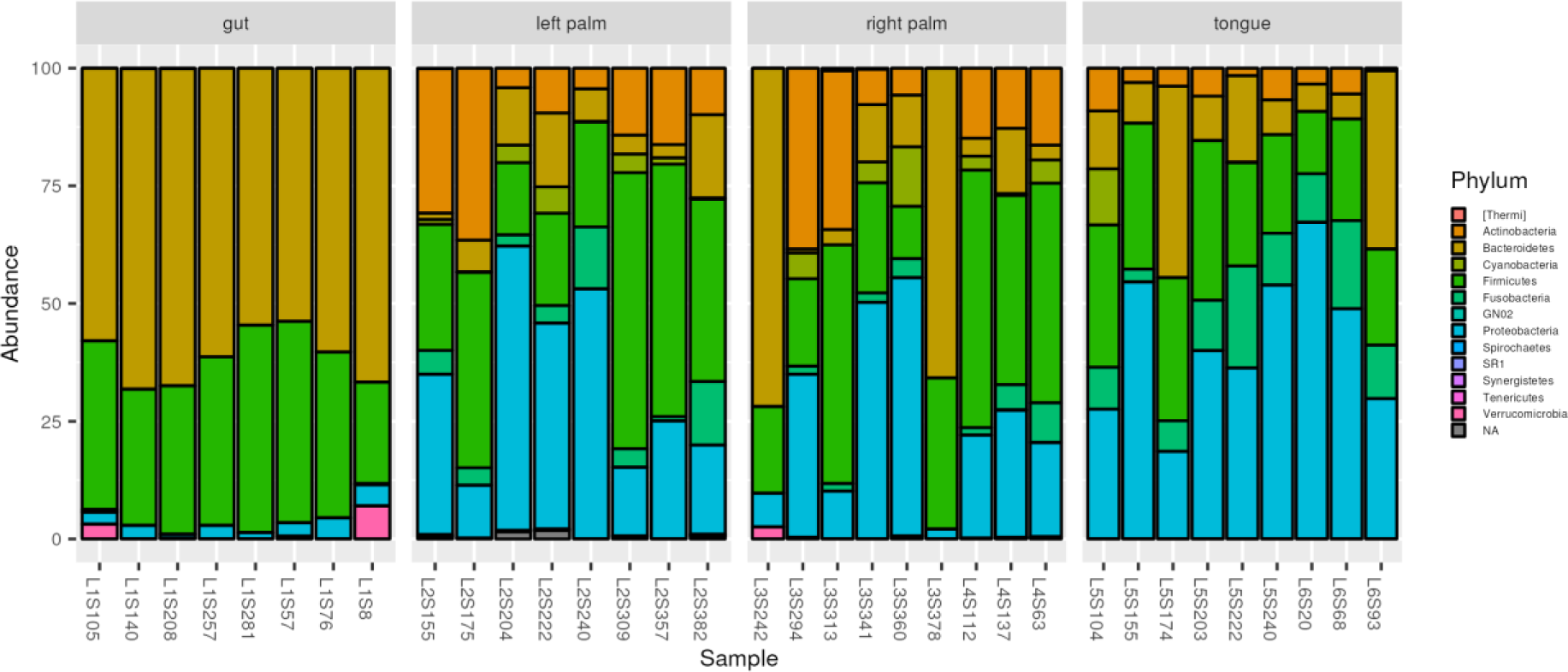
Microbial Abundance by Group: A comparative visualization of the microbial community composition across different body sites, showcasing the diversity and relative abundance of microbial taxa.

Fig 2 reveals distinct microbial community compositions across different body sites, emphasizing the site-specific nature of the human microbiome. This suggests that microbial communities are adapted to the unique environmental conditions of each body site, which could have implications for understanding site-specific microbial functions and their impact on human health [16].

**Fig 2.**
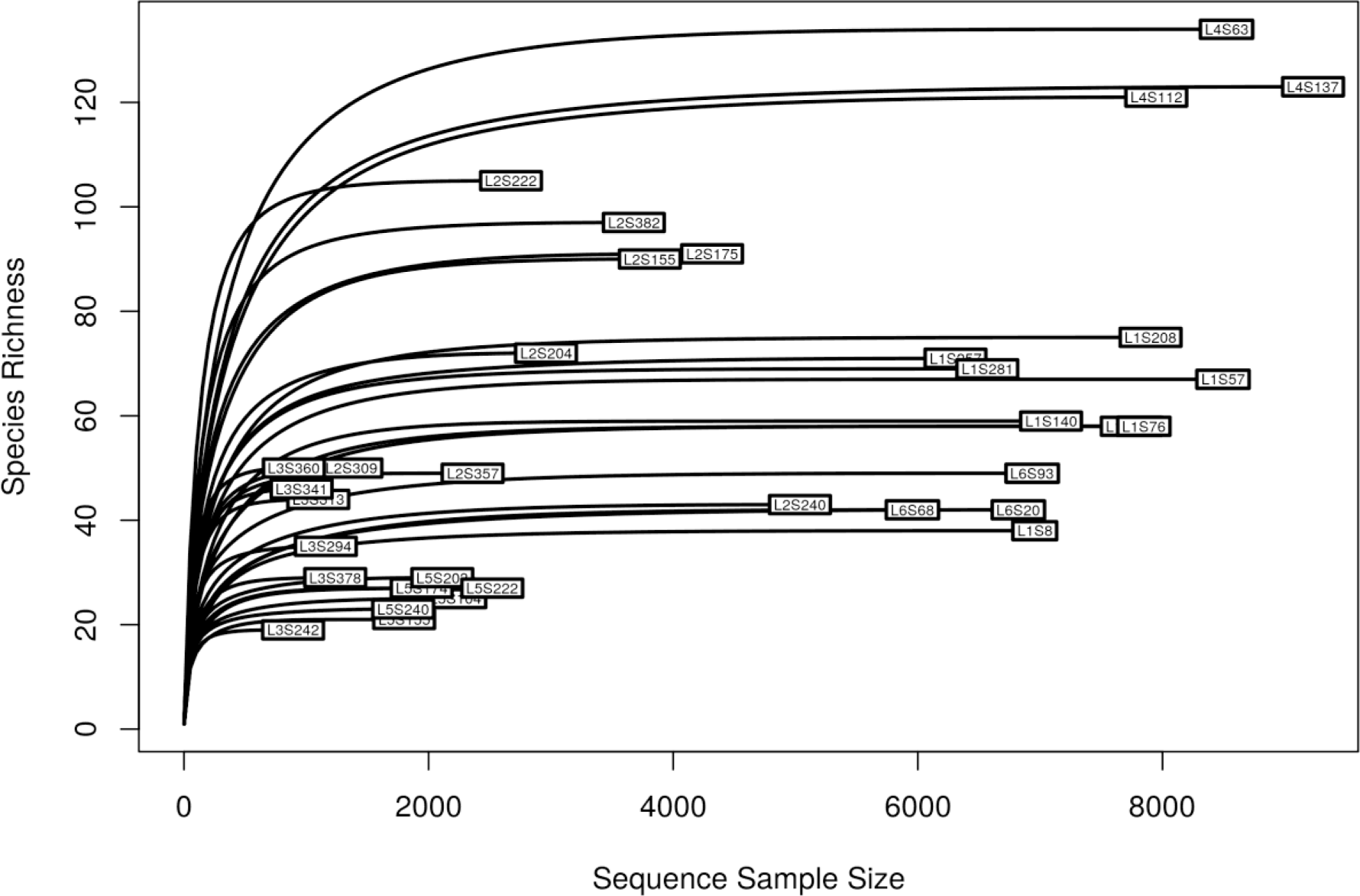
Alpha Rarefaction Curve: This plot illustrates the relationship between sampling depth and observed microbial diversity within samples. Each curve represents a different sample, showing how species richness (alpha diversity) increases with the number of sequences sampled. The plateau indicates the point at which additional sequencing yields minimal new diversity, suggesting adequate sampling depth for capturing the community’s complexity.

The human microbiome, the collection of trillions of microbes living in and on the human body, plays a crucial role in maintaining health and influencing disease. The composition of these microbial communities varies significantly across different body sites, such as the gut, skin, oral cavity, and urogenital tract, reflecting the adaptation of microbial taxa to the local environmental conditions of each site [17,28]. This site-specific nature of the microbiome is a fundamental aspect of human microbial ecology, with important implications for understanding the interactions between microbes and their host, as well as for the development of targeted therapeutic interventions.

The distinct compositions observed in Fig 2 underscore the concept that microbial communities are finely tuned to the specific conditions of their local environment. Factors such as pH, temperature, moisture, and the presence of specific host-derived nutrients and antimicrobial substances contribute to shaping the unique microbial ecosystem of each body site [28,29]. For instance, the skin microbiome is influenced by factors like sebum production and skin moisture, leading to the predominance of lipid-tolerant bacteria in sebaceous areas [30]. Similarly, the gut microbiome is shaped by diet, gut motility, and the immune system, resulting in a diverse community dominated by anaerobic bacteria capable of fermenting dietary fibers [31].

Understanding the site-specific nature of the human microbiome has profound implications for human health. It is well-established that disruptions in the normal microbiome, known as dysbiosis, can contribute to a wide range of diseases, including inflammatory bowel disease, obesity, and skin conditions like eczema [32,33]. Moreover, the site-specific microbiome plays a critical role in protecting against pathogen colonization, modulating the immune system, and contributing to the metabolism of drugs and dietary compounds [34,35].

### Abundance Boxplots per Feature

Boxplots detailing the abundance of microbial features across samples highlighted the variability and distribution of key taxa. This analysis aids in identifying features that are consistently present across samples or those that show significant variation.

The variability in microbial feature abundance across samples, as shown in Fig 3, indicates a diverse microbial presence across the human body. This diversity within and between individuals highlights the complexity of the human microbiome and the need for large-scale studies to capture the full extent of microbial diversity [17].

**Fig 3.**
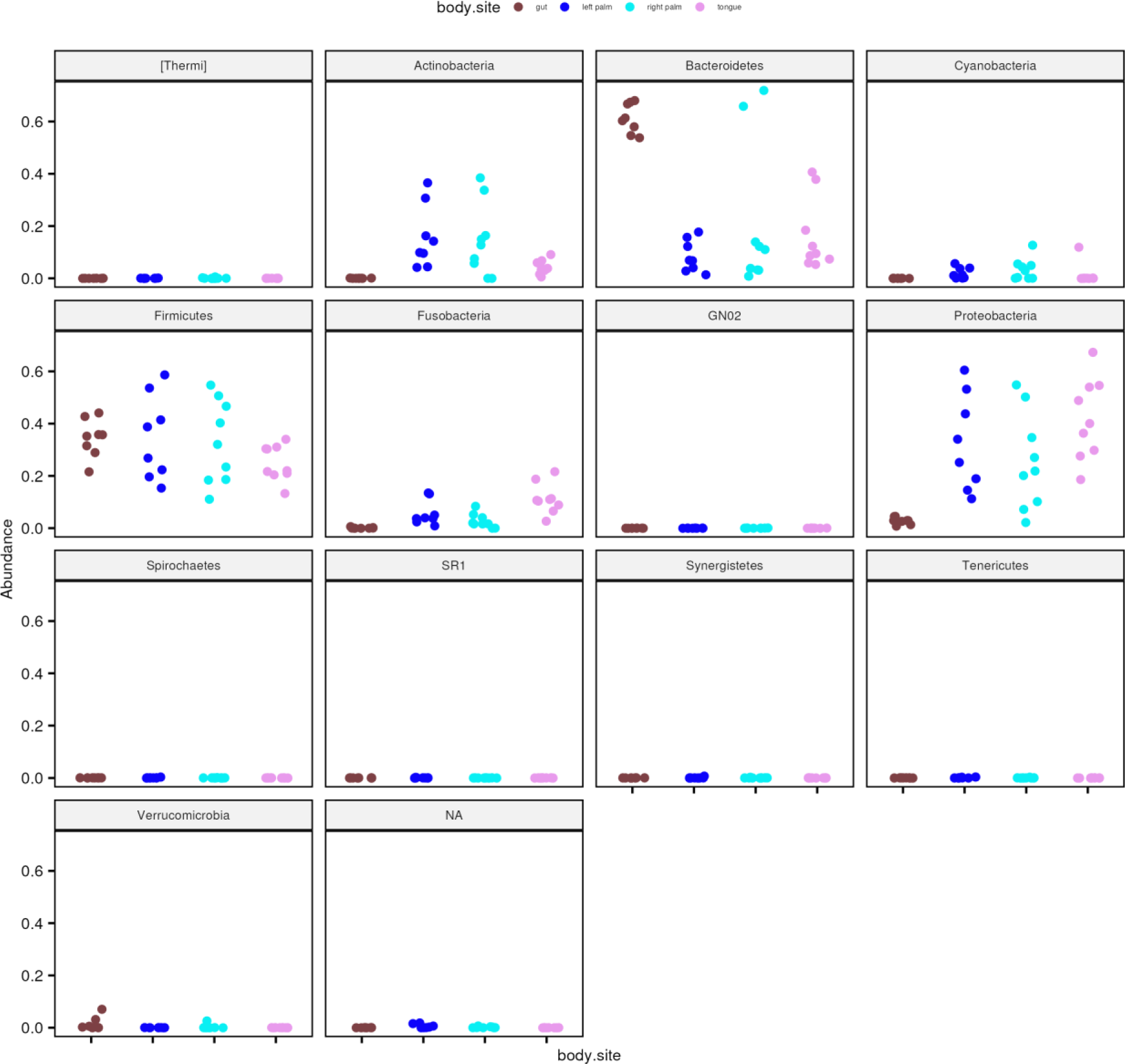
Variability in Microbial Feature Abundance: Boxplots illustrating the distribution and variability of microbial features across all samples, providing insights into the commonality and rarity of specific taxa.

Boxplots, as depicted in Fig 3, are a powerful statistical tool used to visualize the distribution and variability of microbial features across samples. These plots provide a clear visual representation of the central tendency, dispersion, and outliers within the data, making them invaluable for microbiome research.

Features that are consistently present across samples, often referred to as core microbiota, play fundamental roles in maintaining host health by contributing to metabolic processes, protecting against pathogens, and modulating the immune system [36]. Conversely, taxa that show significant variation among samples may reflect individual differences in lifestyle, genetics, diet, or environmental exposures [37].

The insights gained from analyzing the variability and distribution of microbial features have profound implications for microbiome research. Studies have shown that a diverse microbiome is often associated with good health, whereas reduced diversity has been linked to various diseases, including obesity, diabetes, and inflammatory bowel disease [16,38]. Therefore, understanding the factors that contribute to microbial variability is essential for developing strategies to manipulate the microbiome for therapeutic purposes.

Moreover, the identification of core microbiota and highly variable taxa can inform the design of probiotics and microbiome-based interventions. By targeting microbial features that are consistently present across healthy individuals, researchers can develop strategies to restore or enhance beneficial microbial functions [39].

The analysis of microbial feature abundance through boxplots, as shown in Fig 3, offers critical insights into the commonality and rarity of specific taxa across samples.

### Top Four Taxa Boxplots with Wilcoxon p-values

We conducted a Wilcoxon test to assess the significance of differences between groups for the top four taxa. The resulting boxplots, annotated with p-values, indicate statistically significant differences in the abundance of these taxa across different body sites, offering insights into the microbial composition unique to each site. Particularly, the abundance of feature:

- d29fe3c70564fc0f69f2c03e0d1e5561, classified as “Bacteria| Firmicutes| Bacilli| Lactobacillales| Streptococcaceae| Streptococcus” is significantly different between gut and tongue body sites (p-value 0.018). This finding may reflect the role of Streptococcus species in the gut microbiome, where they contribute to the breakdown of complex carbohydrates and production of lactic acid, influencing gut health and disease [40].
- 4b5eeb300368260019c1fbc7a3c718fc, classified as “Bacteria| Bacteroidetes| Bacteroidia| Bacteroidales| Bacteroidaceae| Bacteroides” is significantly different between gut and left palm (p-value 0.00016), gut and right palm (p-value 0.027), and gut and tongue (p-value 8.2e-05). Bacteroides within the gut environment play a crucial role in fermenting polysaccharides to produce short-chain fatty acids beneficial for colon health [41].
- 1d2e5f3444ca750c85302ceee2473331, classified as “Bacteria| Proteobacteria| Gammaproteobacteria| Pasteurellales| Pasteurellaceae| Haemophilus| parainfluenzae” is significantly different between gut and left palm (p-value 0.0009), gut and right palm (p-value 0.0044), and gut and tongue (p-value 0.00061). This could indicate a niche specialization of H. parainfluenzae, with implications for understanding its role in oral and respiratory health [42].
- a049763053c277b16c2a318f41eb23b4, classified as “Bacteria| Actinobacteria| Actinobacteria| Actinomycetales| Corynebacteriaceae| Corynebacterium” is significantly different between left palm and tongue (p-value 0.0055), gut and tongue (p-value 0.00048), gut and right palm (p-value 0.0095) and gut and left palm (p-value 0.00068). Corynebacterium species are known for their role in skin health, contributing to the skin’s barrier function and potentially influencing skin disorders [43].

The significant differences in the abundance of top taxa across body sites, as demonstrated in Fig 4, suggest that certain microbial taxa are more prevalent or exclusive to specific body sites. This could reflect adaptations of these taxa to local environmental conditions or their roles in site-specific physiological processes. Understanding these differences is crucial for exploring the potential functional implications of these taxa in health and disease. The significant differences in taxa abundance suggest adaptations to local environmental conditions or specific physiological roles, offering potential targets for therapeutic interventions.

**Fig 4.**
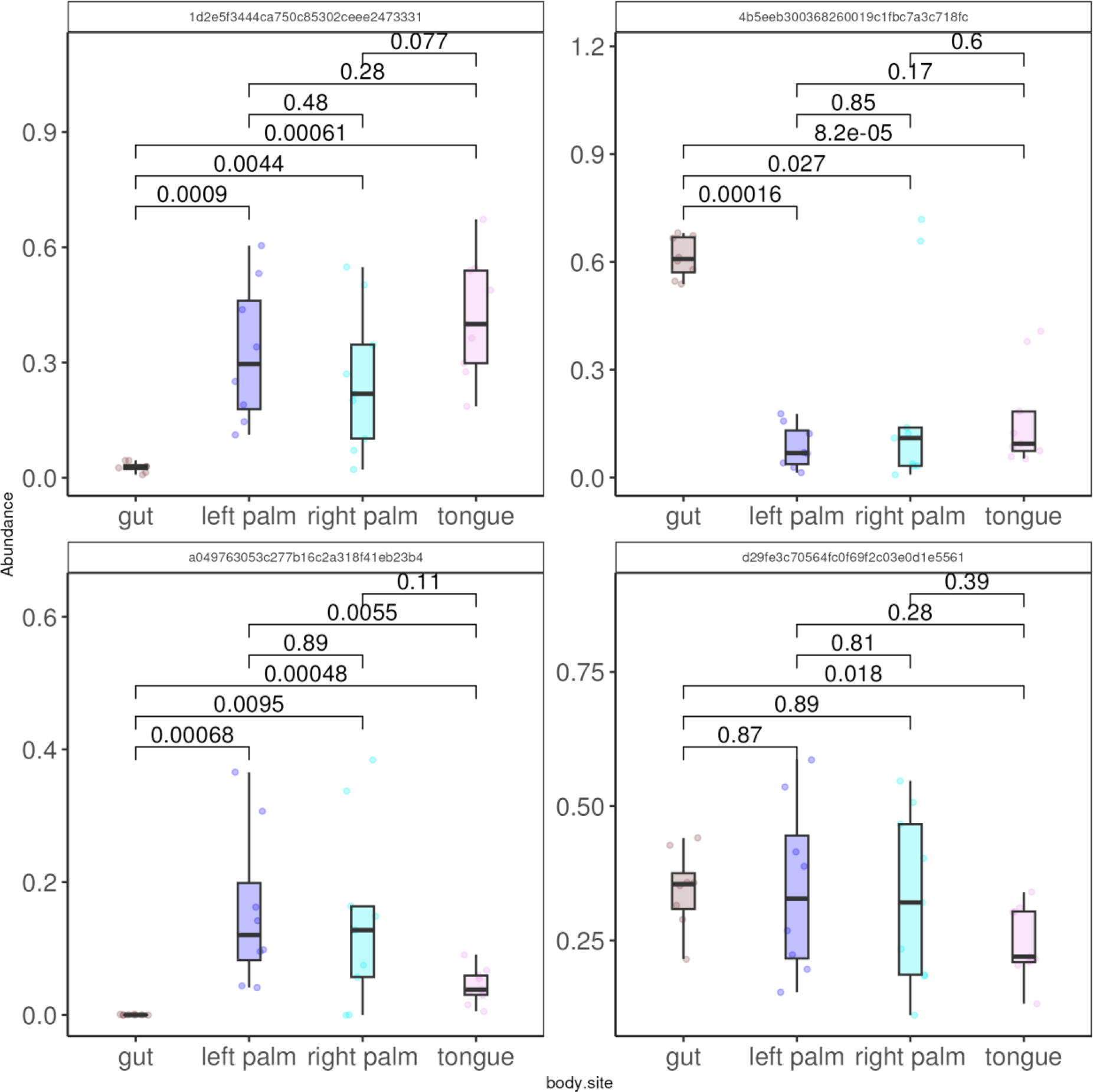
Statistical Comparison of Top Four Taxa Across Groups: Boxplots annotated with Wilcoxon test p-values, highlighting significant differences in the abundance of the top four taxa between different body sites.

### Mean Relative Abundance Plot

The mean relative abundance plot for various taxa across different body sites, along with standard deviations, provides a comprehensive view of the microbial landscape. Notably, the gut is enriched in Bacteroidetes and depleted in Proteobacteria, contrasting with other body sites that show a higher abundance of Proteobacteria. Moreover, right and left palm are enriched in Actinobacteria, while other body sites are not.

Fig 5 provides a clear view of the dominant microbial taxa across different body sites, highlighting the gut’s unique microbial landscape compared to other body sites. This suggests that the gut microbiome may play distinct roles in human health, potentially influencing digestion, immune function, and even behavior.

**Fig 5.**
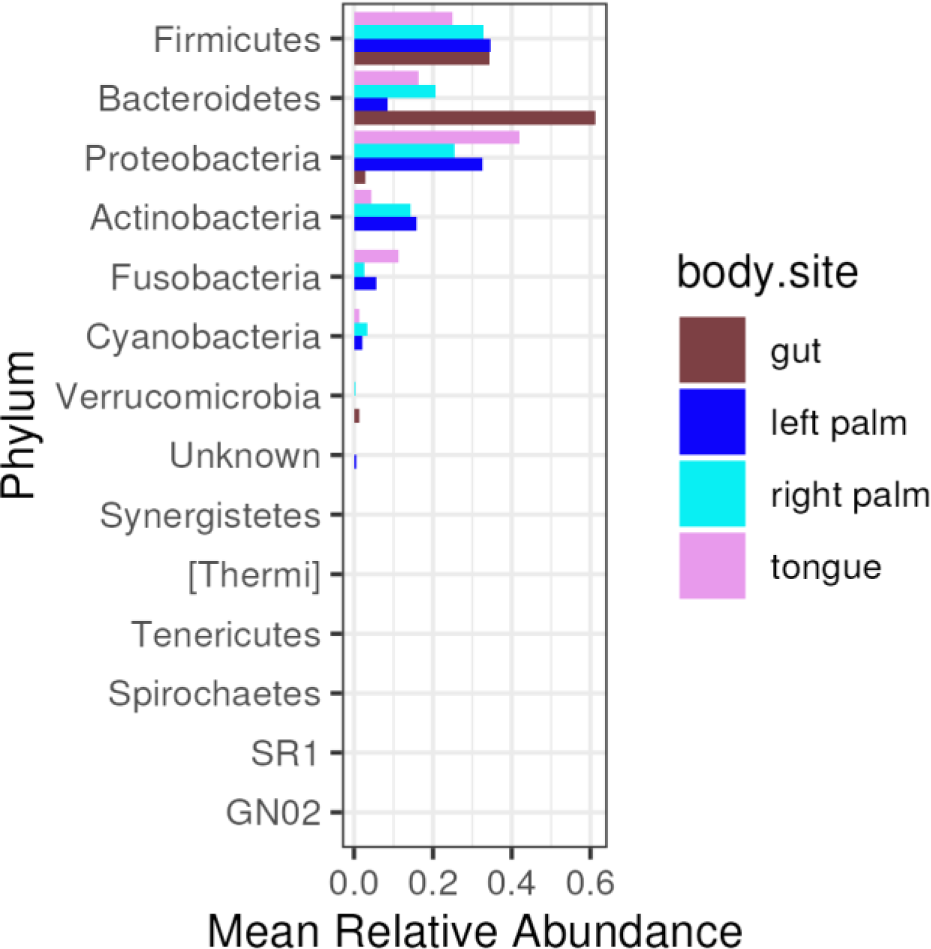
Mean Relative Abundance of Microbial Taxa: A plot detailing the mean relative abundance and standard deviations of various microbial taxa across different body sites, offering a clear view of microbial distribution patterns.

The gut microbiome is particularly interesting due to its dense microbial population and its significant impact on human health. It has been observed that the gut is enriched in Bacteroidetes, a phylum of Gram-negative bacteria, and depleted in Proteobacteria, another phylum that includes many pathogenic bacteria such as Escherichia, Salmonella, and Helicobacter. Bacteroidetes are known for their ability to degrade complex polysaccharides, which is crucial for the digestion of dietary fiber [41]. This capability supports the host’s energy metabolism and is linked to maintaining a healthy body weight and preventing metabolic disorders [33].

In contrast, a higher abundance of Proteobacteria in other body sites, or even in the gut, is often associated with dysbiosis and has been linked to various diseases, including inflammatory bowel disease (IBD) and obesity [44]. The enrichment of Bacteroidetes in the gut and the relative depletion of Proteobacteria may, therefore, reflect a balance conducive to health.

The skin microbiome, particularly on the palms, shows a distinct composition compared to other body sites. Actinobacteria, especially the genus Corynebacterium, are enriched on the skin of the palms. Actinobacteria are known for their role in skin health, contributing to the skin’s barrier function and producing antimicrobial compounds that protect against pathogens [43]. The enrichment of Actinobacteria on the palms may be related to the specific environmental conditions of the skin, such as dryness and exposure to the external environment, which favor the growth of these bacteria.

### Dominant taxa

The gut body site is dominated by Bacteroidetes phylum, with a relative frequency percentage of 100% across samples. The tongue body site is dominated by Proteobacteria phylum, with a prevalence frequency percentage of 67%, followed by Bacteroidetes 22% and Firmicutes 11%. The left palm is dominated 50%-50% by phyla Proteobacteria and Firmicutes. The right palm is dominated by Firmicutes 44%, Bacteroidetes 22% and Proteobacteria 22% and Actinobacteria 11%.

The Table 1 shows mean relative abundances and standard deviations for all groups:

**Table 1.**
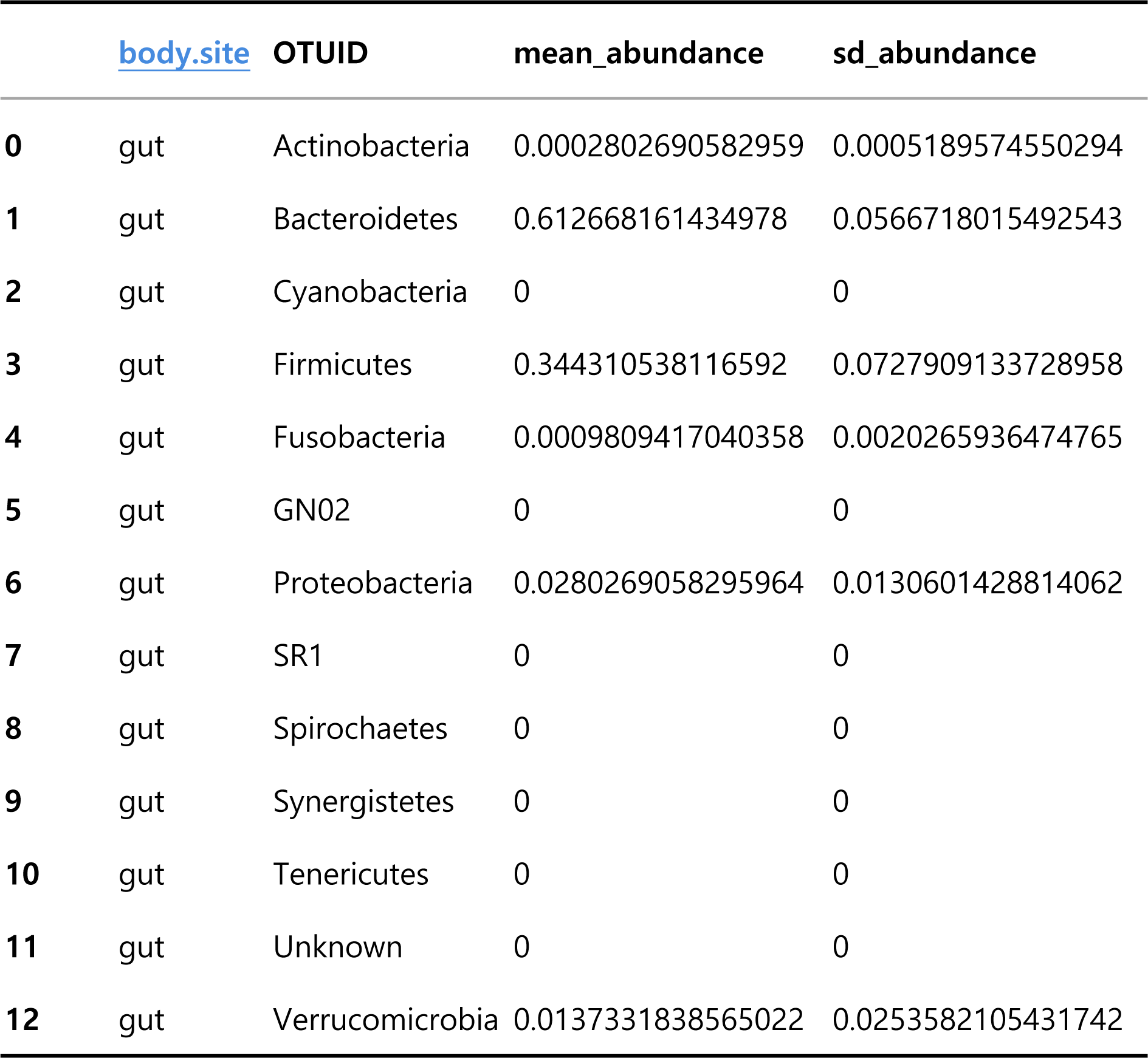
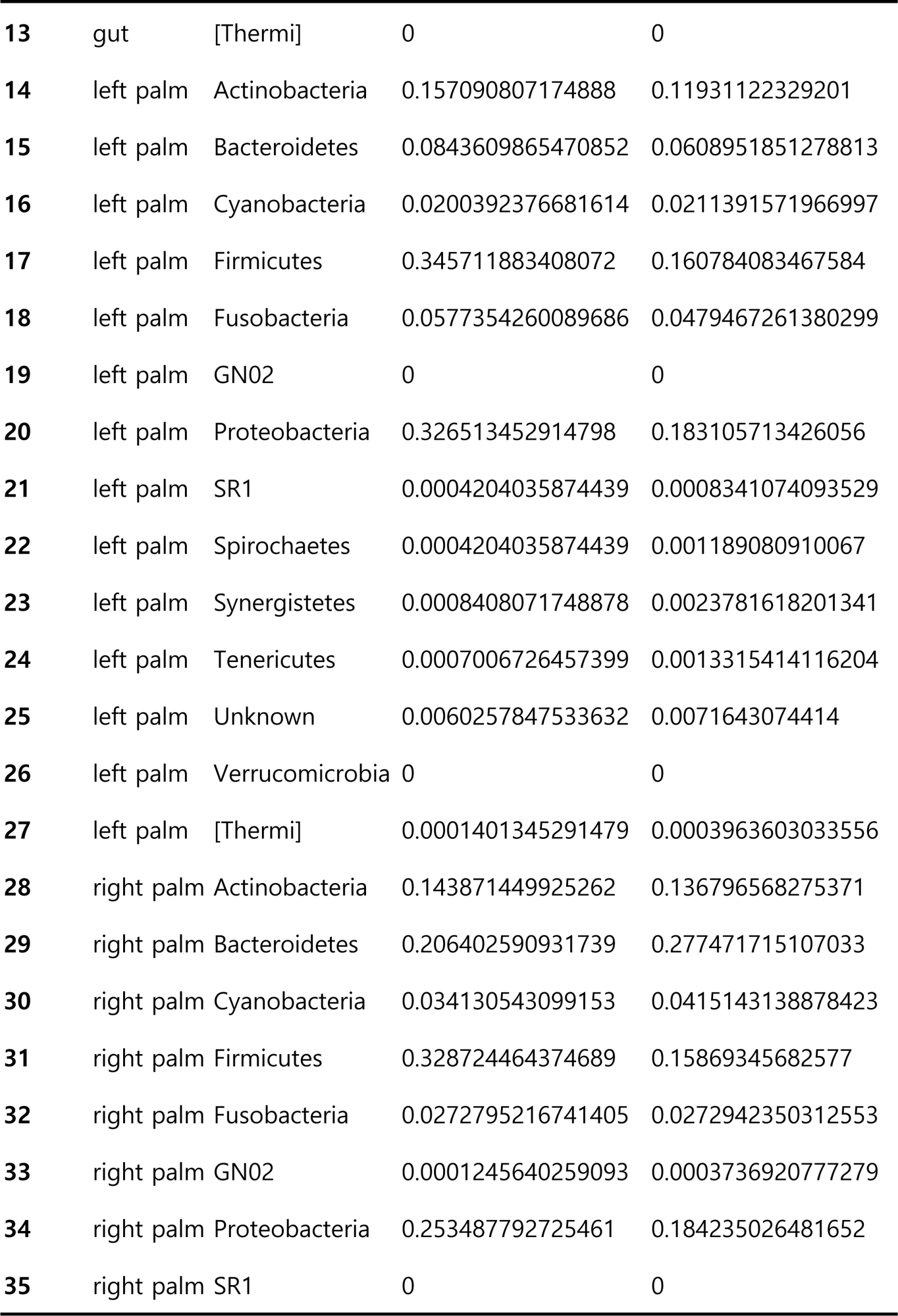
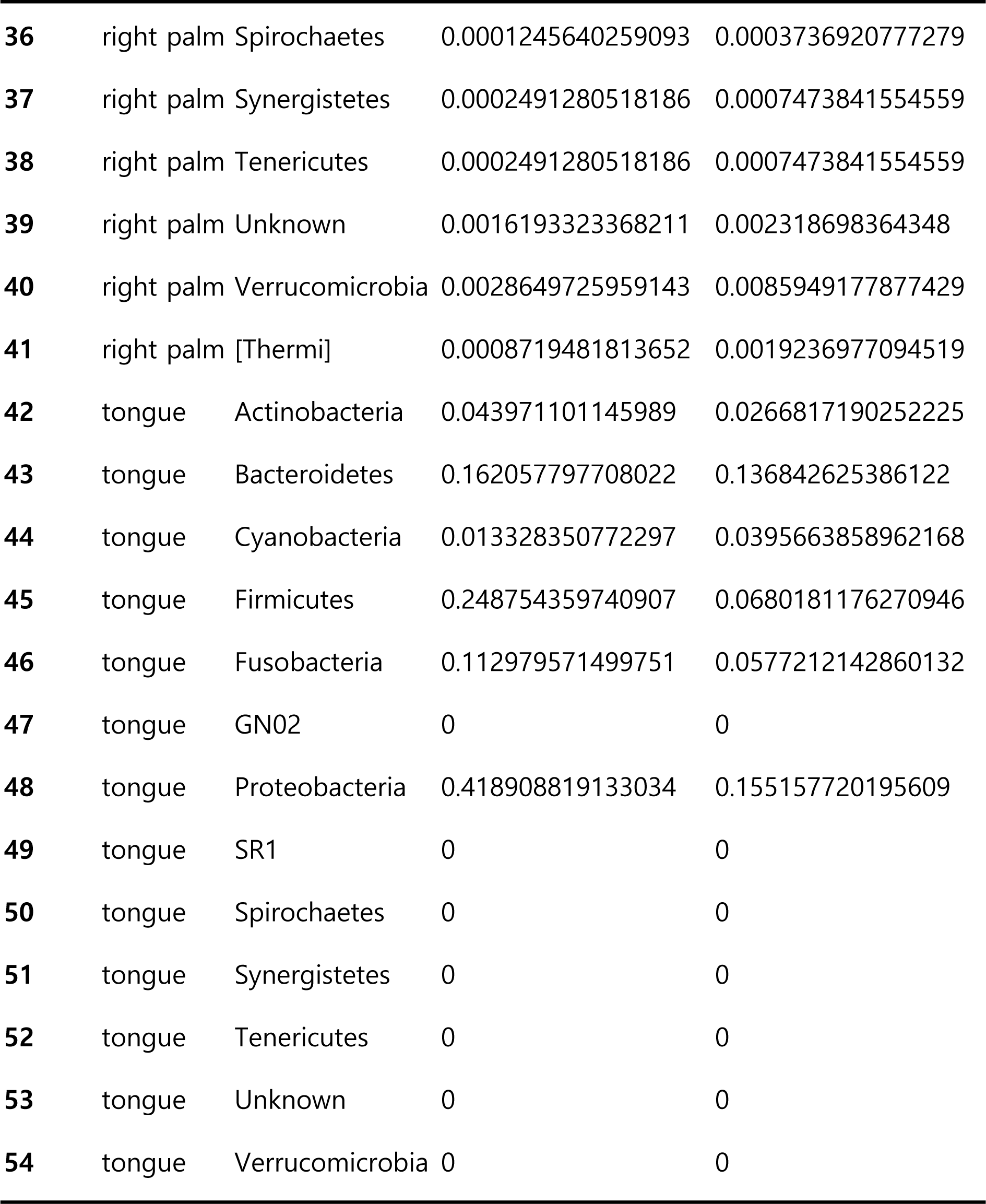
Mean relative abundances and standard deviations of features.

|  | body.site | OTUID | mean_abundance | sd_abundance |
| --- | --- | --- | --- | --- |
| 0 | gut | Actinobacteria | 0.0002802690582959 | 0.0005189574550294 |
| 1 | gut | Bacteroidetes | 0.612668161434978 | 0.0566718015492543 |
| 2 | gut | Cyanobacteria | 0 | 0 |
| 3 | gut | Firmicutes | 0.344310538116592 | 0.0727909133728958 |
| 4 | gut | Fusobacteria | 0.0009809417040358 | 0.0020265936474765 |
| 5 | gut | GN02 | 0 | 0 |
| 6 | gut | Proteobacteria | 0.0280269058295964 | 0.0130601428814062 |
| 7 | gut | SR1 | 0 | 0 |
| 8 | gut | Spirochaetes | 0 | 0 |
| 9 | gut | Synergistetes | 0 | 0 |
| 10 | gut | Tenericutes | 0 | 0 |
| 11 | gut | Unknown | 0 | 0 |
| 12 | gut | Verrucomicrobia | 0.0137331838565022 | 0.0253582105431742 |
| **13** | gut | [Thermi] | 0 | 0 |
| **14** | left palm | Actinobacteria | 0.157090807174888 | 0.11931122329201 |
| **15** | left palm | Bacteroidetes | 0.0843609865470852 | 0.0608951851278813 |
| **16** | left palm | Cyanobacteria | 0.0200392376681614 | 0.0211391571966997 |
| **17** | left palm | Firmicutes | 0.345711883408072 | 0.160784083467584 |
| **18** | left palm | Fusobacteria | 0.0577354260089686 | 0.0479467261380299 |
| **19** | left palm | GN02 | 0 | 0 |
| **20** | left palm | Proteobacteria | 0.326513452914798 | 0.183105713426056 |
| **21** | left palm | SR1 | 0.0004204035874439 | 0.0008341074093529 |
| **22** | left palm | Spirochaetes | 0.0004204035874439 | 0.001189080910067 |
| **23** | left palm | Synergistetes | 0.0008408071748878 | 0.0023781618201341 |
| **24** | left palm | Tenericutes | 0.0007006726457399 | 0.0013315414116204 |
| **25** | left palm | Unknown | 0.0060257847533632 | 0.0071643074414 |
| **26** | left palm | Verrucomicrobia | 0 | 0 |
| **27** | left palm | [Thermi] | 0.0001401345291479 | 0.0003963603033556 |
| **28** | right palm | Actinobacteria | 0.143871449925262 | 0.136796568275371 |
| **29** | right palm | Bacteroidetes | 0.206402590931739 | 0.277471715107033 |
| **30** | right palm | Cyanobacteria | 0.034130543099153 | 0.0415143138878423 |
| **31** | right palm | Firmicutes | 0.328724464374689 | 0.15869345682577 |
| **32** | right palm | Fusobacteria | 0.0272795216741405 | 0.0272942350312553 |
| **33** | right palm | GN02 | 0.0001245640259093 | 0.0003736920777279 |
| **34** | right palm | Proteobacteria | 0.253487792725461 | 0.184235026481652 |
| **35** | right palm | SR1 | 0 | 0 |
| 36 | right palm | Spirochaetes | 0.0001245640259093 | 0.0003736920777279 |
| 37 | right palm | Synergistetes | 0.0002491280518186 | 0.0007473841554559 |
| 38 | right palm | Tenericutes | 0.0002491280518186 | 0.0007473841554559 |
| 39 | right palm | Unknown | 0.0016193323368211 | 0.002318698364348 |
| 40 | right palm | Verrucomicrobia | 0.0028649725959143 | 0.0085949177877429 |
| 41 | right palm | [Thermi] | 0.0008719481813652 | 0.0019236977094519 |
| 42 | tongue | Actinobacteria | 0.043971101145989 | 0.0266817190252225 |
| 43 | tongue | Bacteroidetes | 0.162057797708022 | 0.136842625386122 |
| 44 | tongue | Cyanobacteria | 0.013328350772297 | 0.0395663858962168 |
| 45 | tongue | Firmicutes | 0.248754359740907 | 0.0680181176270946 |
| 46 | tongue | Fusobacteria | 0.112979571499751 | 0.0577212142860132 |
| 47 | tongue | GN02 | 0 | 0 |
| 48 | tongue | Proteobacteria | 0.418908819133034 | 0.155157720195609 |
| 49 | tongue | SR1 | 0 | 0 |
| 50 | tongue | Spirochaetes | 0 | 0 |
| 51 | tongue | Synergistetes | 0 | 0 |
| 52 | tongue | Tenericutes | 0 | 0 |
| 53 | tongue | Unknown | 0 | 0 |
| 54 | tongue | Verrucomicrobia | 0 | 0 |

### Correlation Analysis

A heatmap of the Spearman correlation between the top 30 Amplicon Sequence Variants (ASVs) and continuous metadata variables, transformed with Centered Log-Ratio (CLR), reveals patterns of association between microbial features and clinical biomarkers. The Centered Log-Ratio (CLR) transformation is applied to the data before conducting the correlation analysis to address the compositional nature of microbiome data. Microbiome datasets are inherently compositional since they represent relative abundances of microbial taxa within samples, summing up to a constant total [45]. The CLR transformation, a standard approach in compositional data analysis, mitigates the issues arising from the ‘constant sum constraint’ by converting the data into log-ratio space, facilitating more meaningful statistical comparisons and correlations [46].

This analysis underscores the potential implications of microbial composition for health and disease. In the specific case under study, no meaningful conclusions can be extrapolated from this kind of analysis, and it is solely reported to demonstrate the capabilities of MicrobiomePhylo. The test for significant spearman correlations, with p-values adjusted using Benjamini-Hochberg correction, reported no significant correlations found.

The heatmap in Fig 6, although not yielding significant correlations in this study, illustrates the potential of correlation analysis in uncovering relationships between microbial features and clinical biomarkers. This approach could reveal how changes in microbial composition relate to health outcomes, guiding future microbiome research and therapeutic interventions.

**Fig 6.**
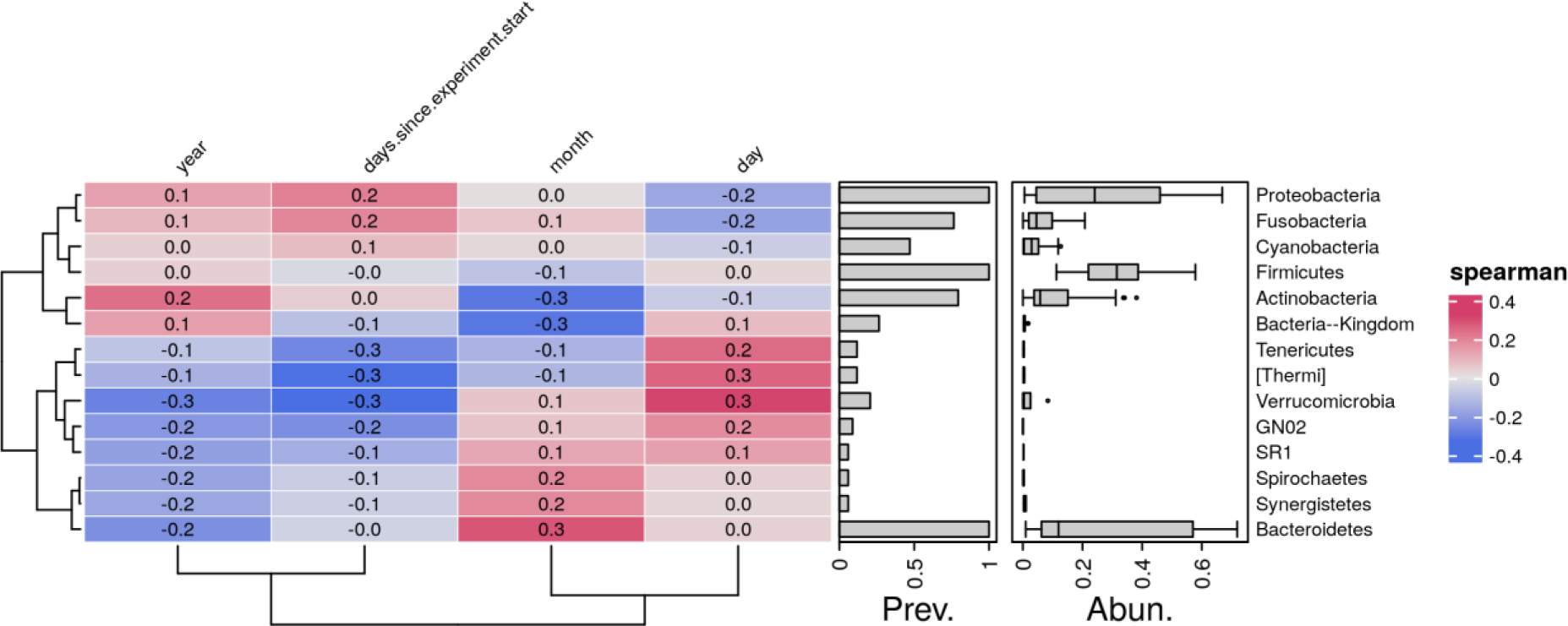
Correlation Analysis of Microbial Features and Metadata: Heatmap displaying the Spearman correlation between the top 30 Amplicon Sequence Variants (ASVs) and continuous metadata variables, transformed with CLR and clustered for enhanced interpretability.

### Heatmap with Clustering of Samples and Features

Our heatmap analysis, incorporating hierarchical clustering [47], offers a detailed visualization of microbial community structure within the dataset. This approach facilitates the identification of patterns and similarities among samples and features. The heatmap in Fig 7, with its hierarchical clustering of samples and features, reveals a distinct pattern of gut enrichment in Firmicutes and Bacteroidetes phyla compared to other body sites. These two phyla are known to dominate the human gut microbiome, playing crucial roles in metabolism and immune function [48].

**Fig 7.**
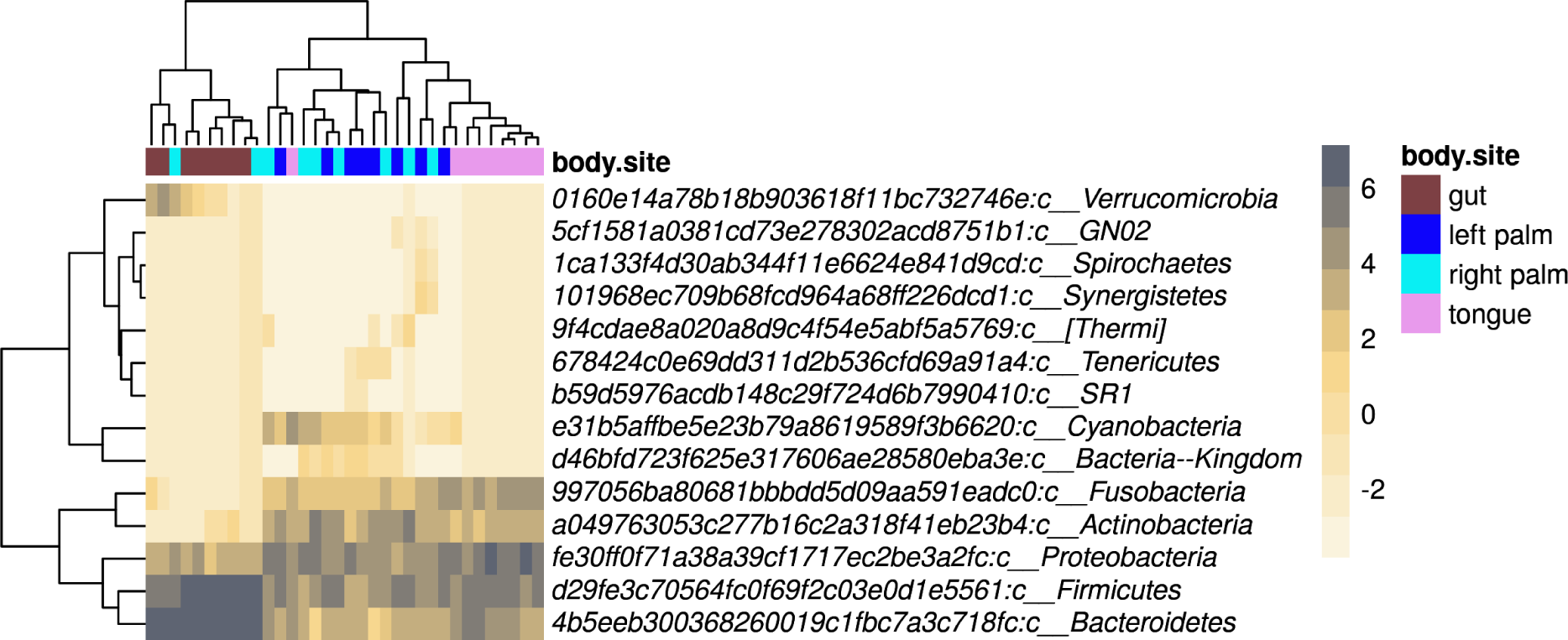
Microbial Community Structure: A comprehensive heatmap with hierarchical clustering of samples and features, revealing patterns of microbial abundance and community composition.

Fig 7’s detailed visualization of microbial community structure within the dataset through hierarchical clustering provides insights into the similarities and differences among samples and features. This could help identify microbial signatures associated with specific body sites or health conditions, offering a basis for targeted microbiome interventions.

### Heatmap per Group with Abundances of Features

Fig 8 emphasizes the variations in microbial community composition associated with different body sites, reinforcing the concept of a site-specific microbiome. Understanding these variations is essential for developing targeted microbiome-based therapies and diagnostics.

**Fig 8.**
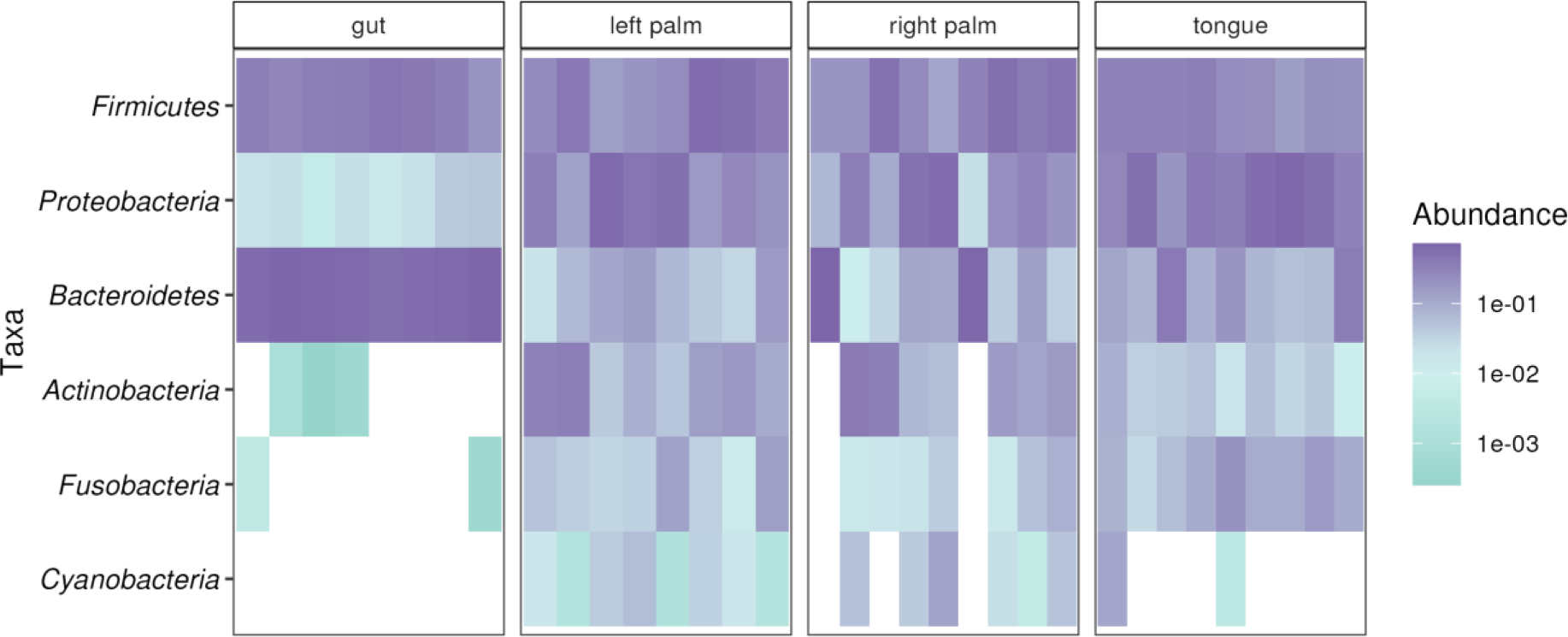
Group-Specific Microbial Community Composition: Heatmaps visualizing the relative abundance of microbial taxa at the specified taxonomic level across different sample groups, highlighting variations associated with the grouping variable.

### Boxplots for All Alpha Diversity Metrics for Each Group

Boxplots representing alpha diversity metrics for each group underscore the diversity within samples.

- Observed Features is a simple count of the unique operational taxonomic units (OTUs), species, or amplicon sequence variants (ASVs) detected in each sample. It is a direct measure of species richness but does not account for the abundance of each species.
- Shannon Entropy (also known as the Shannon Diversity Index) combines both richness and evenness into a single metric. It quantifies the uncertainty in predicting the species identity of an individual randomly selected from a sample. Higher values indicate greater diversity, reflecting not only a larger number of species but also a more equitable distribution of species abundances.
- Simpson Index is another measure that accounts for both richness and evenness, but it emphasizes the dominance or concentration of a few species within the community. The index ranges from 0 to 1, where values closer to 0 indicate higher diversity and values closer to 1 suggest a community dominated by one or a few species.
- Chao1 is an estimator of species richness that is particularly useful for accounting for unseen species. It is based on the number of rare species (e.g., singletons and doubletons) within a sample and provides an estimate of the total species richness, including species that were not observed due to sampling limitations.

The alpha diversity metrics represented in Fig 9 reflect the within-sample diversity, highlighting differences in microbial richness and evenness across body sites. These differences could have implications for the resilience of microbial communities to perturbations and their ability to perform essential functions.

**Fig 9.**
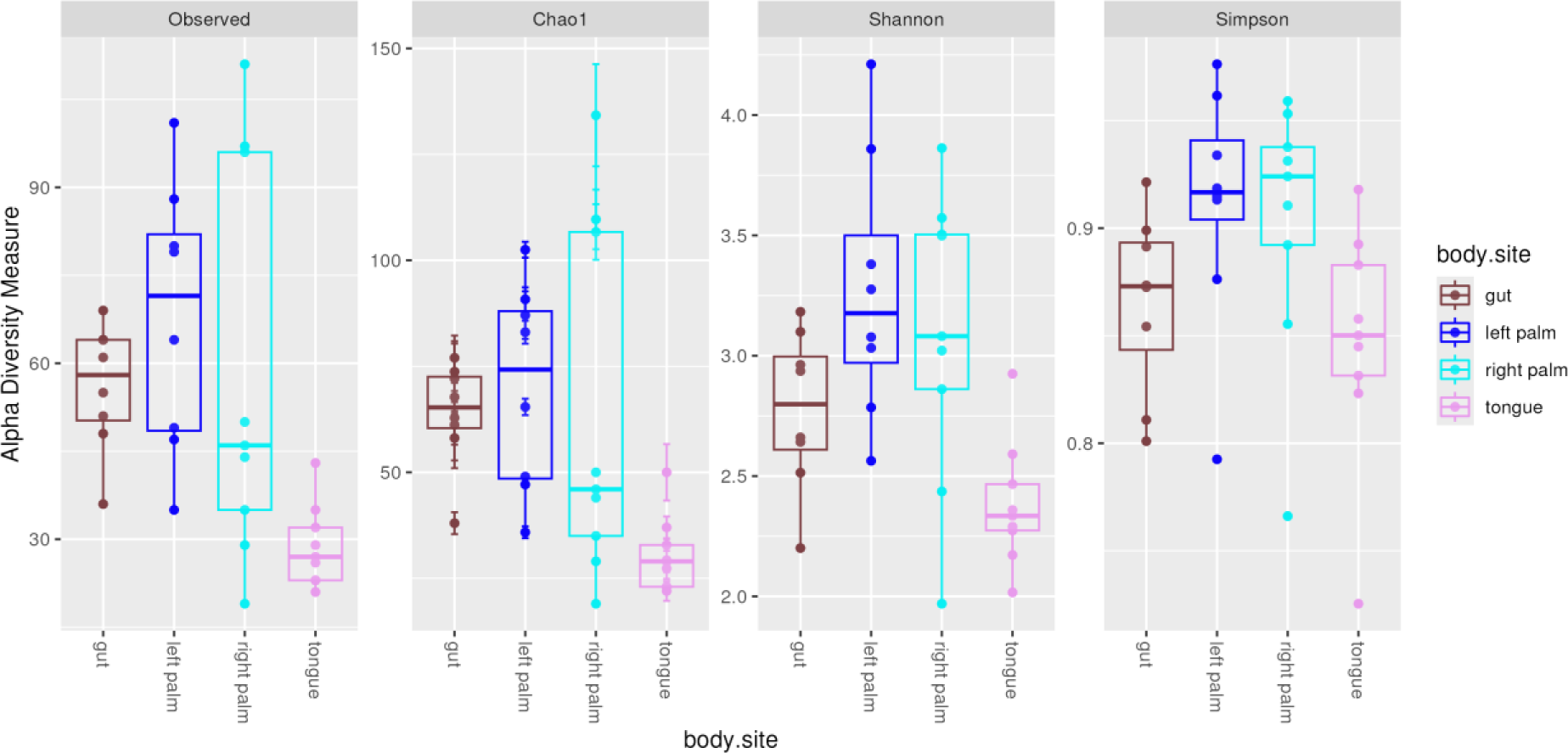
Alpha Diversity Across Groups: Boxplots representing the distribution of alpha diversity metrics (Observed Features, Shannon Entropy, Simpson Index, Chao1) for each group, illustrating within-sample diversity.

For instance, higher diversity (richness and evenness) is often associated with greater stability and resilience to environmental perturbations [38, 49]. Diverse microbial communities are also thought to perform a wider range of functions, contributing to the health and well-being of the host [50].

Moreover, variations in alpha diversity across body sites reflect the distinct ecological niches and selective pressures present in each environment. For example, the gut microbiome typically exhibits higher diversity compared to the skin or oral microbiomes, reflecting the complex interactions and functions required for digestion and immune modulation [16].

### Boxplots Annotated with Kruskal Wallis and Dunn Test Significance for All Alpha Metrics

Kruskal-Wallis tests were performed to identify differences in alpha diversity metrics across groups, followed by Dunn tests for post-hoc analysis.

Effect sizes were calculated to quantify the magnitude of observed differences. Notably, significant differences in Observed Species were observed between gut and tongue (Dunn test p-value 0.0122) and left palm and tongue (p-value 0.0023). Significant differences in Chao1 were observed between gut and tongue (p-value 0.0143) and left palm and tongue (p-value 0.0077). Significant differences in Simpson diversity were not observed. Finally, significant differences in Shannon diversity were observed between the tongue and both palm sites (right palm p-value 0.0216, left palm p-value 0.0047), highlighting distinct microbial communities.

The significant differences in alpha diversity metrics between body sites, as shown in Figs 10-13, suggest that microbial diversity is influenced by the local environment of each body site.

**Fig 10.**
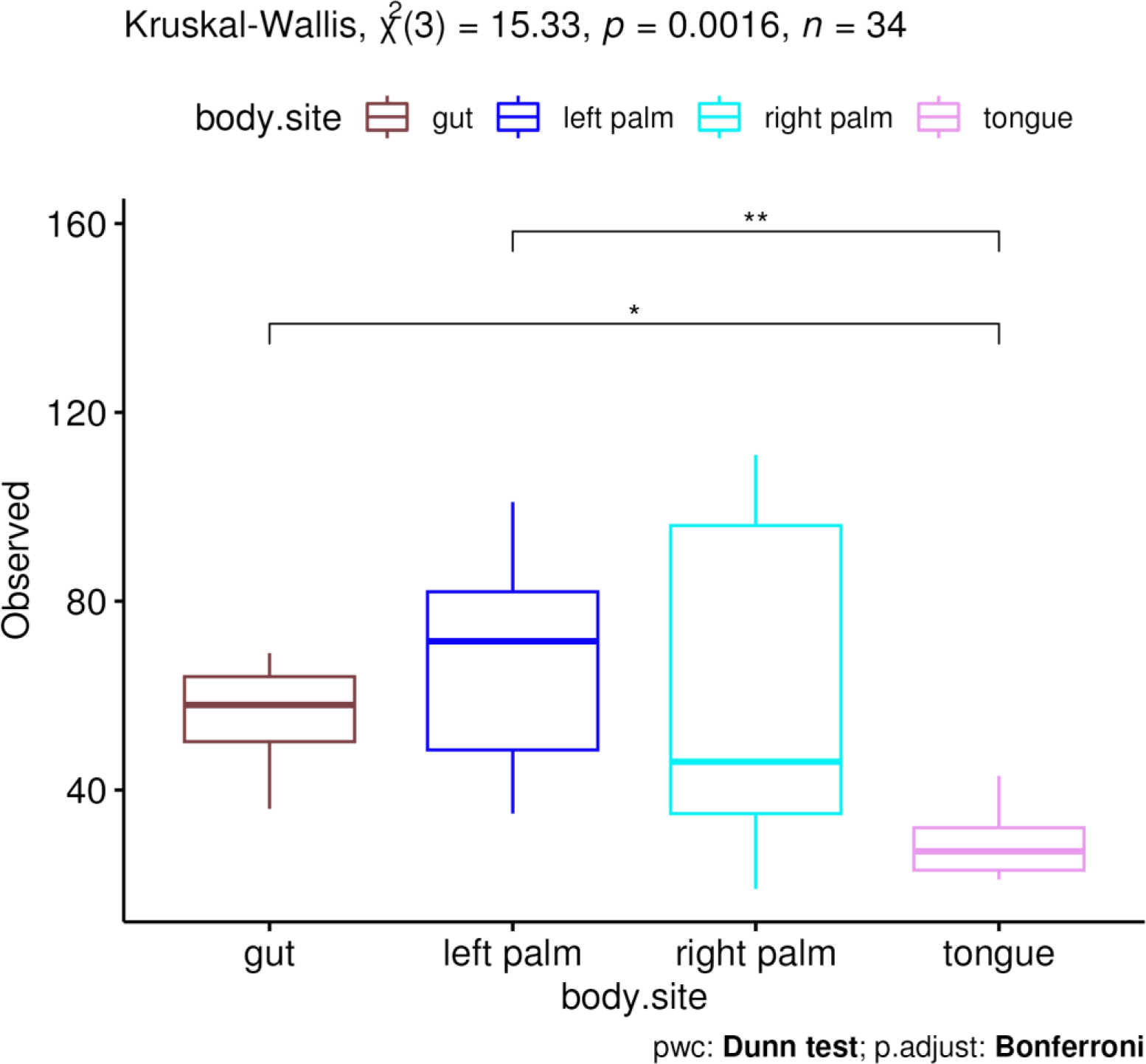
Statistical Analysis of Observed Species: Boxplots annotated with significance levels from Kruskal Wallis and Dunn tests, showcasing significant differences in alpha diversity metrics between groups.

**Fig 11.**
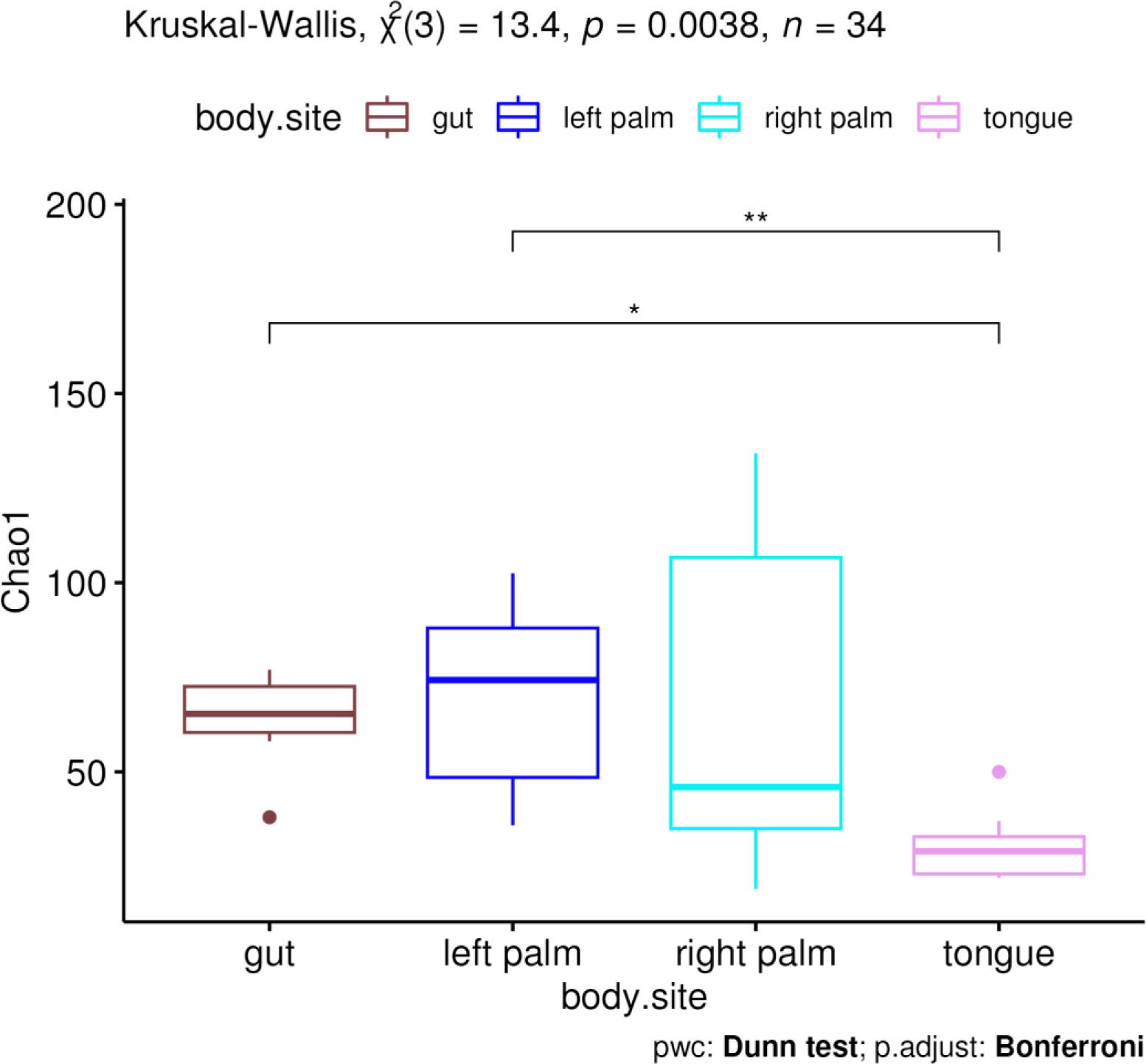
Statistical Analysis of Chao1: Boxplots annotated with significance levels from Kruskal Wallis and Dunn tests, showcasing significant differences in alpha diversity metrics between groups.

**Fig 12.**
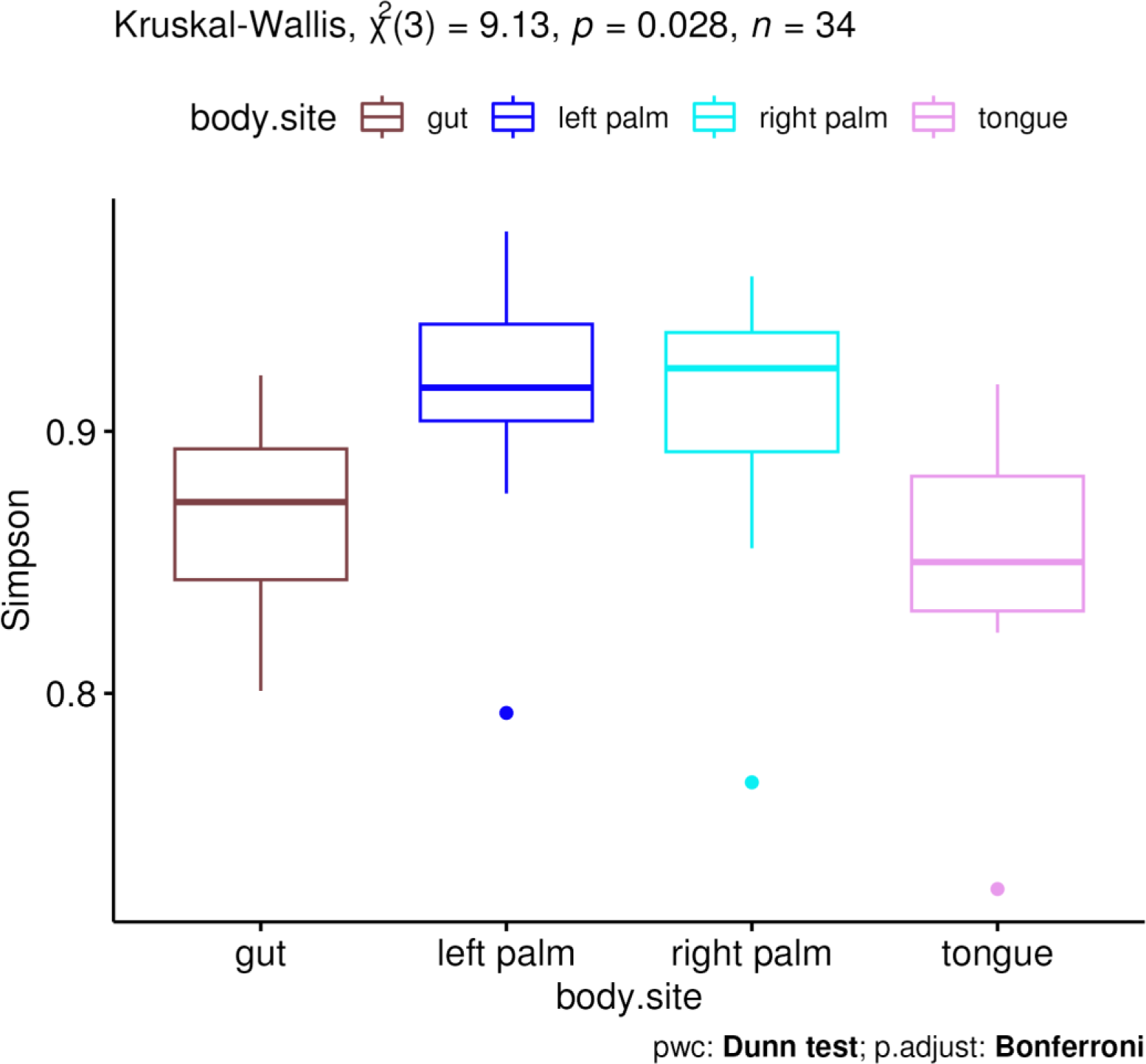
Statistical Analysis of Simpson: Boxplots annotated with significance levels from Kruskal Wallis and Dunn tests, showcasing significant differences in alpha diversity metrics between groups.

**Fig 13.**
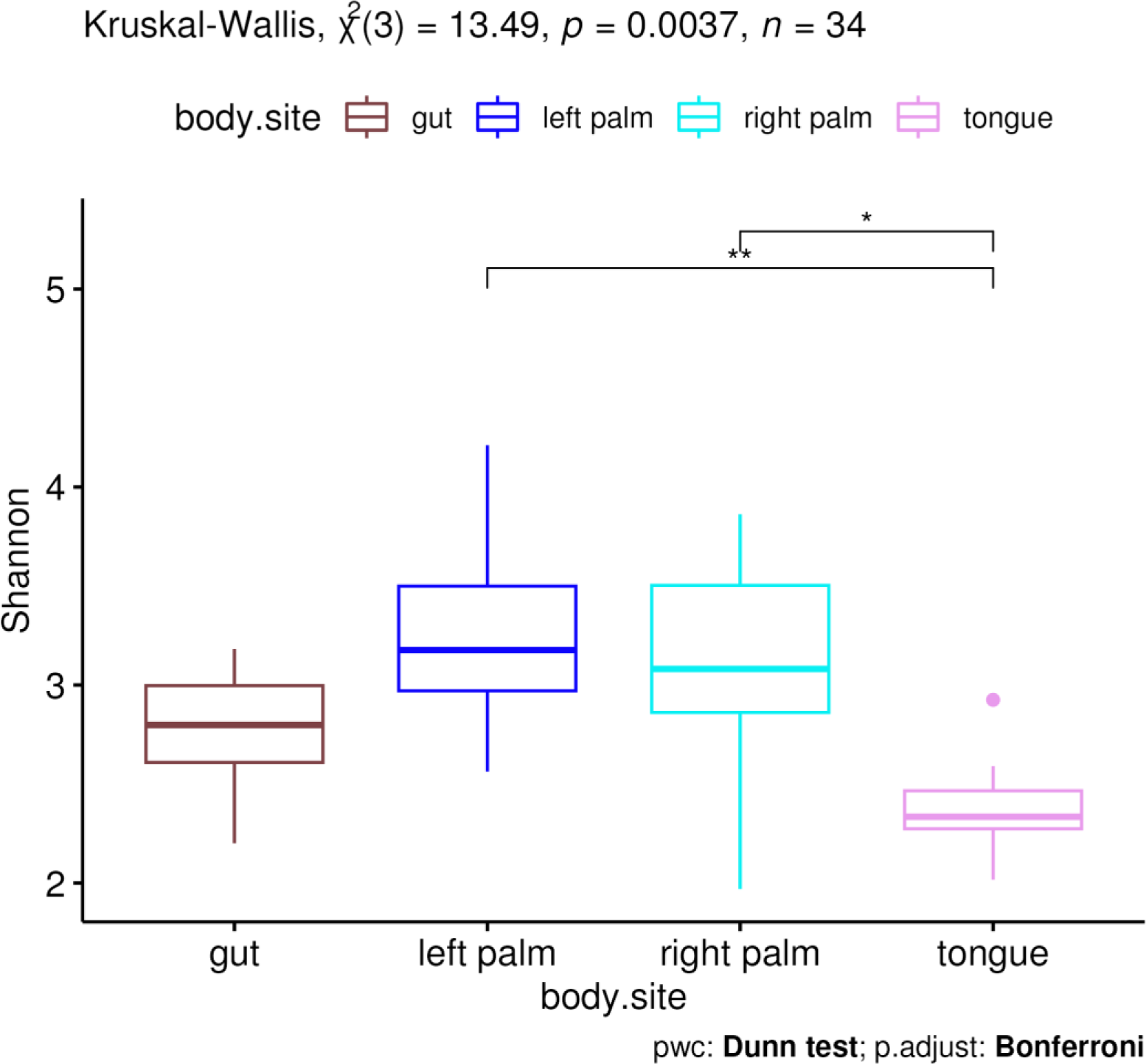
Statistical Analysis of Shannon: Boxplots annotated with significance levels from Kruskal Wallis and Dunn tests, showcasing significant differences in alpha diversity metrics between groups.

These findings are consistent with previous research indicating that the human body hosts distinct microbial communities across different anatomical locations, influenced by factors such as moisture, temperature, pH, and host immune responses [17][28].

The absence of significant differences in Simpson diversity suggests that while the richness and evenness of species may vary between some body sites, the dominance or concentration of a few species does not significantly differ across the sites examined. Conversely, significant differences in Shannon diversity between the tongue and palm sites indicate variations in both species richness and evenness, reflecting the distinct ecological niches of these body sites. This has potential implications for understanding how diversity impacts health and disease resistance.

Higher microbial diversity has been associated with health benefits, including enhanced resistance to pathogen colonization and a more robust immune response [16]. Conversely, reduced diversity has been linked to various diseases, suggesting that alterations in microbial communities could serve as indicators of health status or disease risk [51].

### NMDS and PCoA for Microbial Community Clustering

Both NMDS and PCoA analyses [19][18][20], based on various distance metrics (Jaccard, Bray Curtis, Unifrac, Weighted Unifrac)[6], demonstrated clear clustering of tongue and gut samples, with mixed groups observed for left and right palm samples. These findings suggest distinct microbial community structures associated with different body sites [17]. NMDS operates by preserving the rank order of distances between samples, making it particularly useful for ecological data that may not adhere to linear assumptions. This method is effective in reducing the dimensionality of complex datasets to visualize patterns of similarity and dissimilarity among microbial communities in a low-dimensional space.

**Fig 14.**
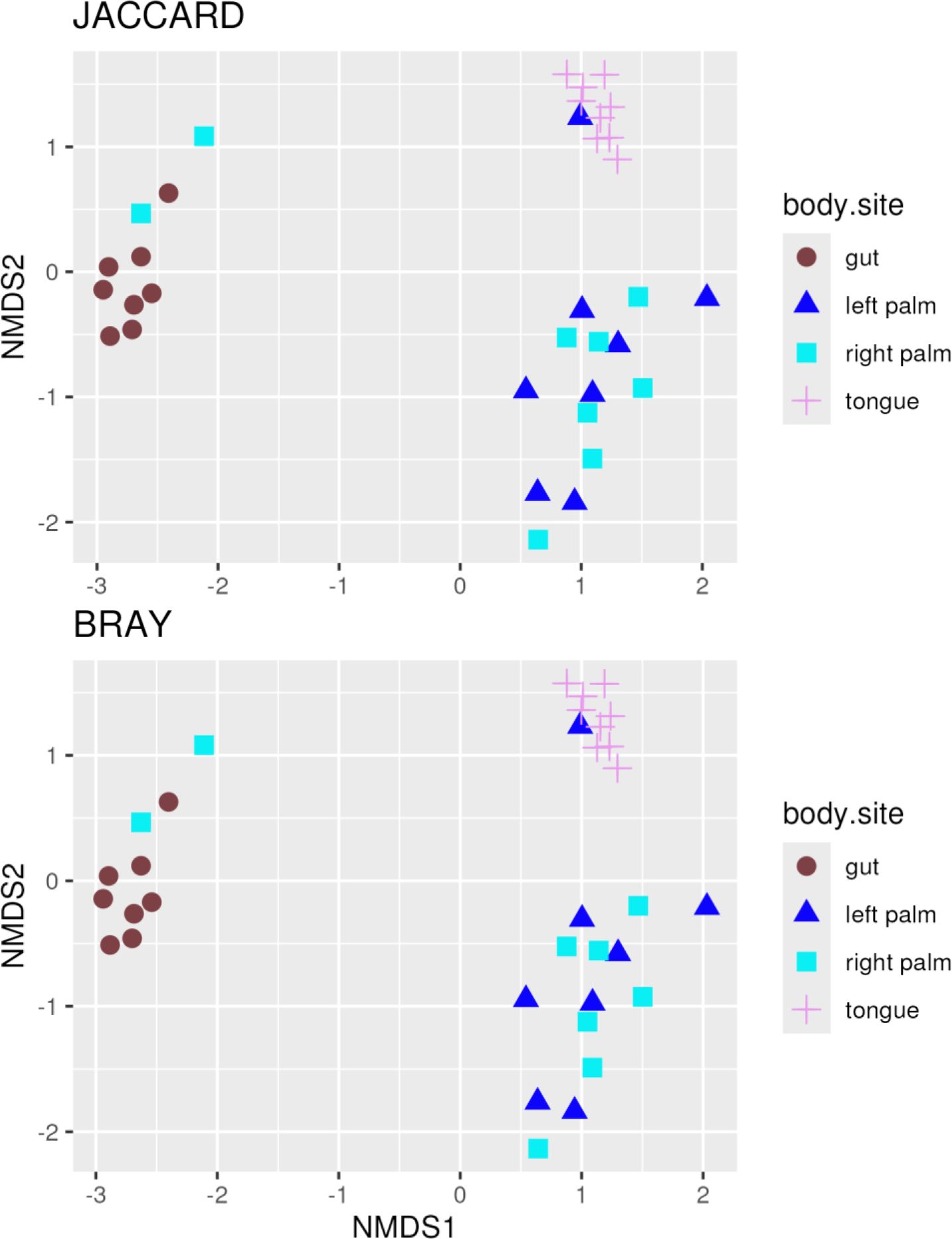
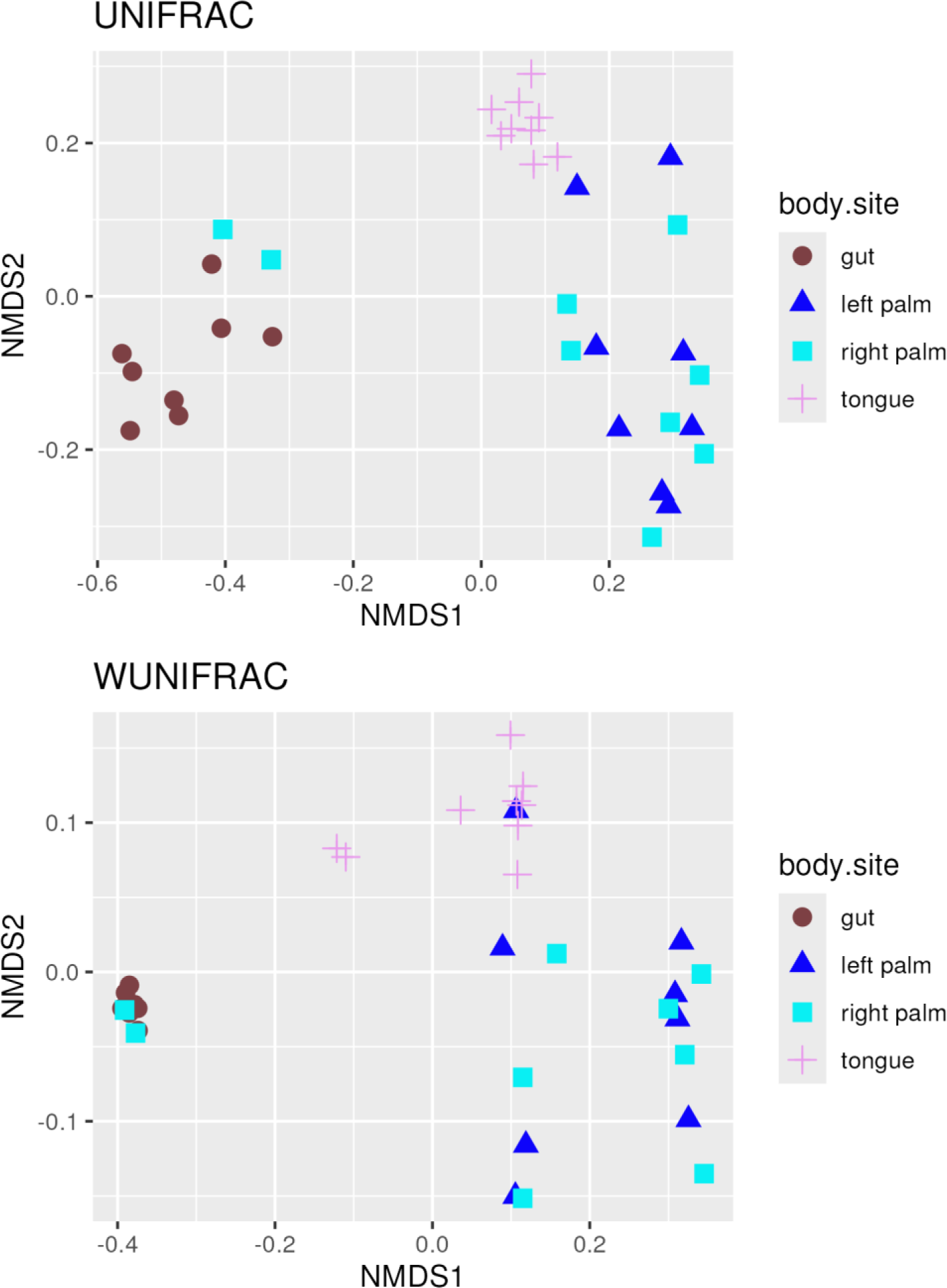
Microbial Community Clustering Analysis: NMDS plots based on Bray Curtis, Jaccard, Unifrac, and Weighted Unifrac distance metrics, demonstrating clear clustering patterns of tongue and gut samples compared to mixed groups of left and right palm.

PCoA, on the other hand, transforms a distance matrix into a new set of orthogonal axes that best represent the variance in the data, allowing for the visualization of microbial community relationships in reduced-dimensional space. The accompanying scree plots for each PCoA analysis (Fig 15) highlight the proportion of variance explained by the principal coordinates, with the first two coordinates typically capturing the most significant patterns of variation among samples. This further supports the distinct clustering patterns of tongue and gut samples, underscoring the unique microbial community structures associated with these body sites.

**Fig 15.**
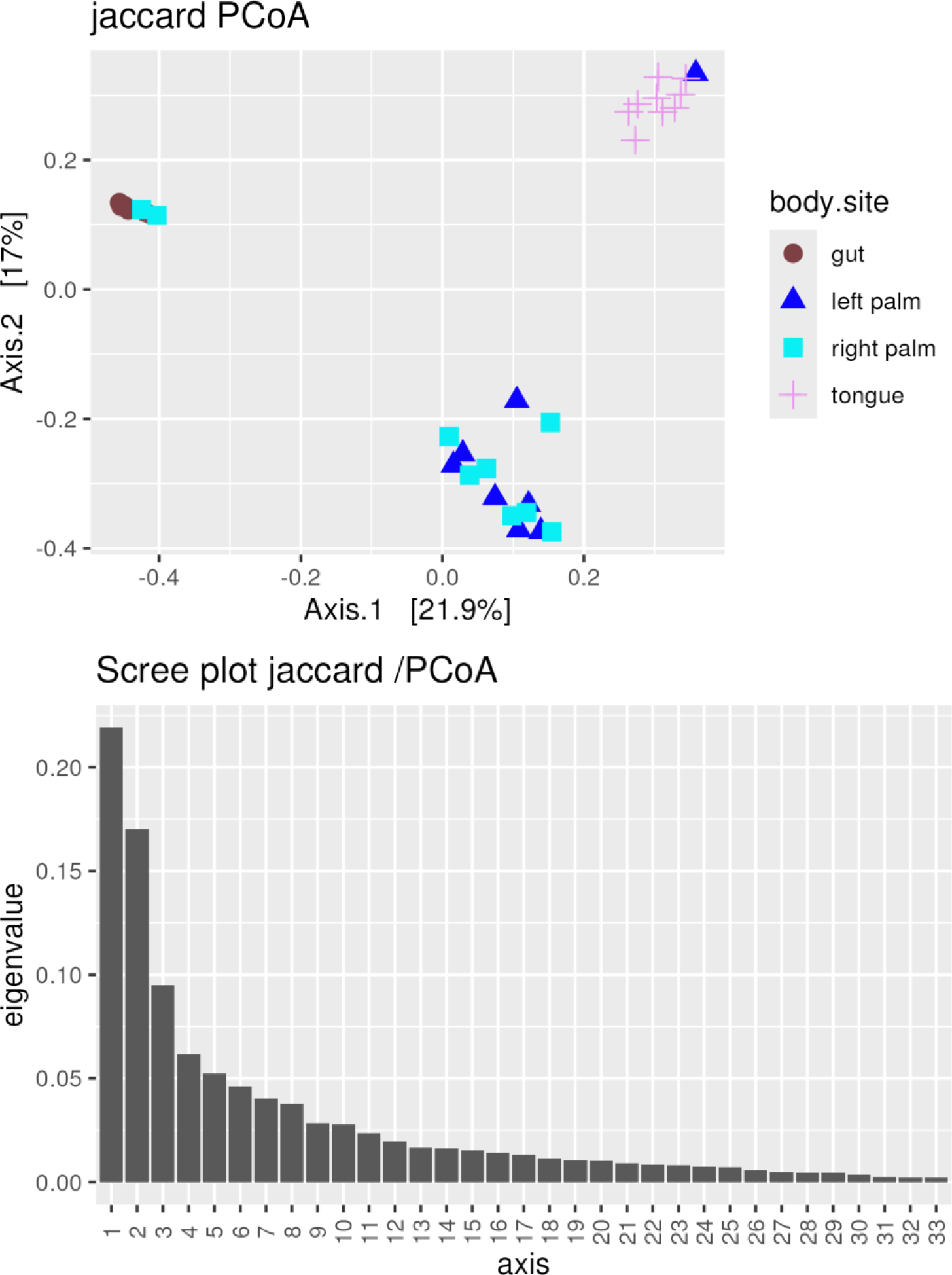
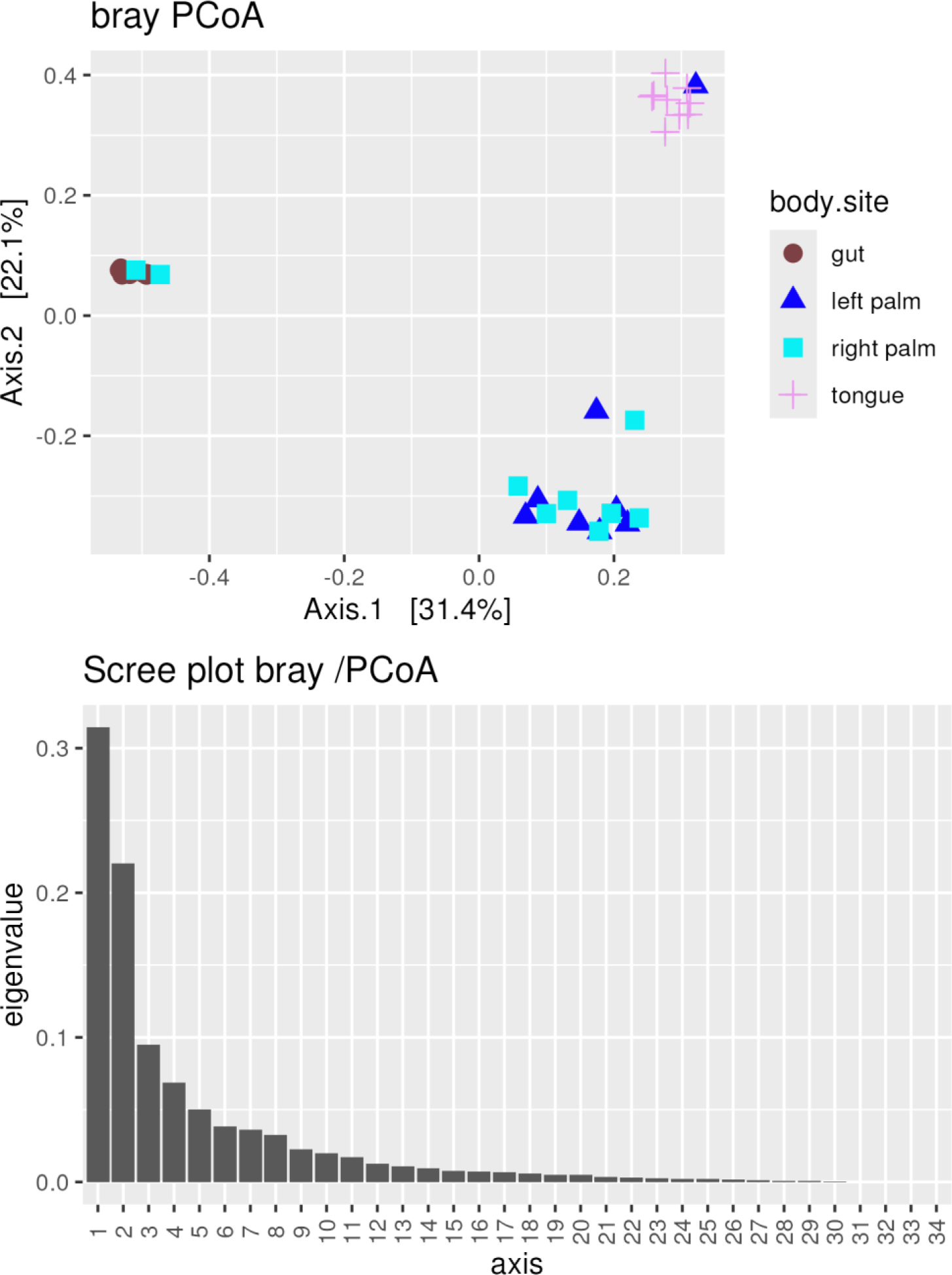
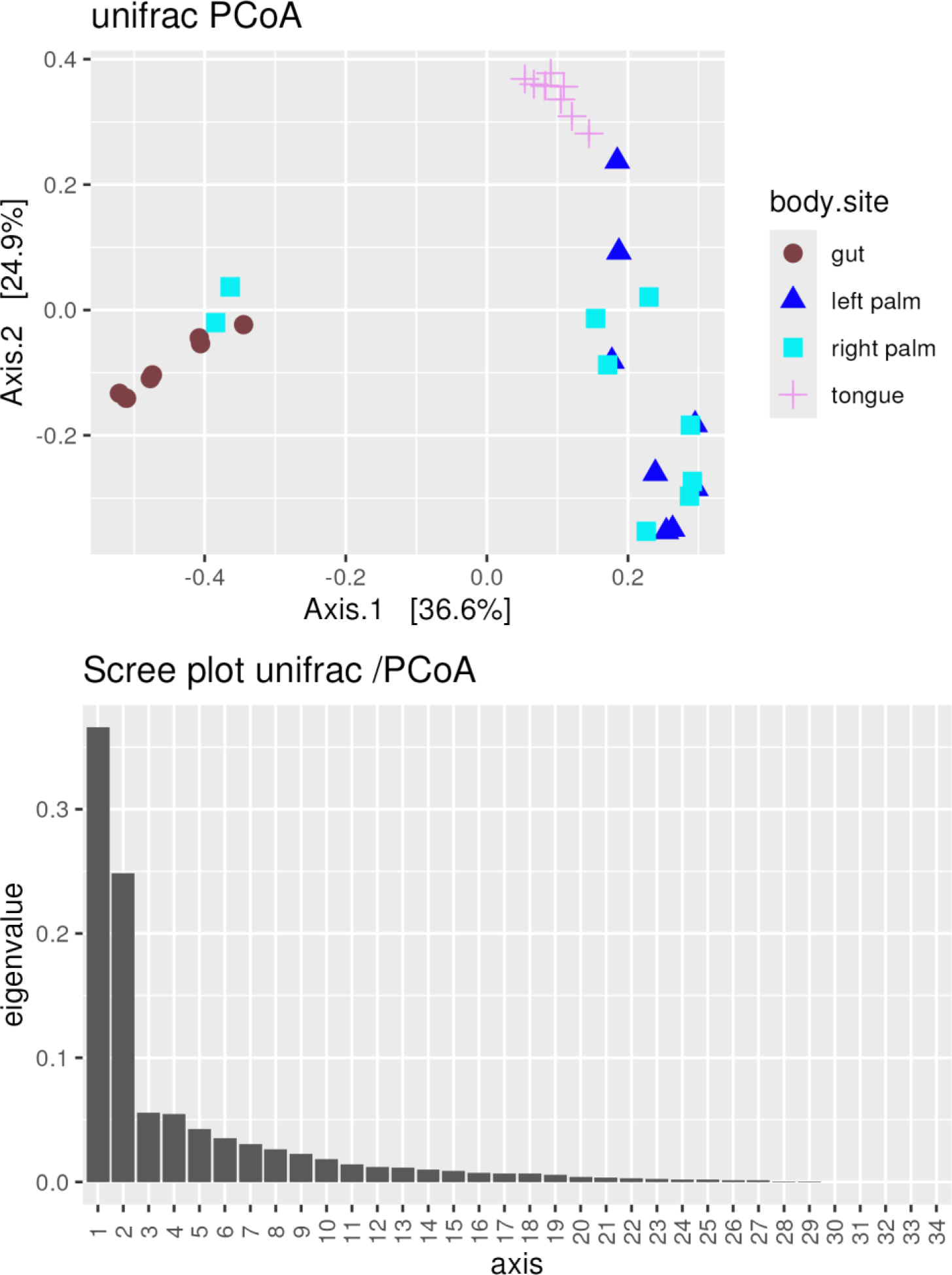
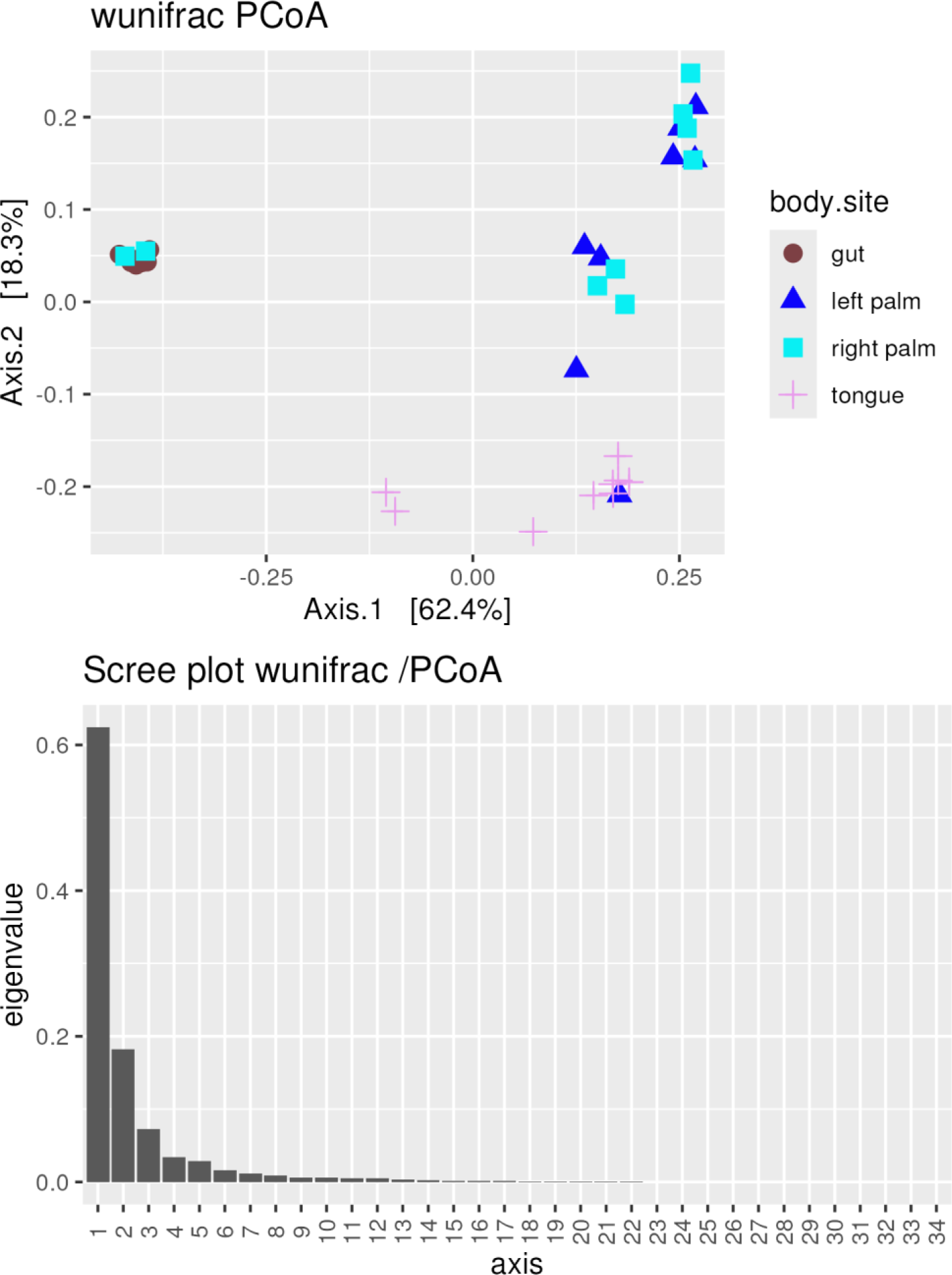
Microbial Community Clustering Analysis: PCoA plots based on Bray Curtis, Jaccard, Unifrac, and Weighted Unifrac distance metrics, demonstrating clear clustering patterns of tongue and gut samples compared to mixed groups of left and right palm. Accompanying scree plots for each PCoA plot, demontrating the high variance explained by the first two principal coordinates for all distance metrics.

The clear clustering of tongue and gut samples in Figs 14 and 15, based on various distance metrics, indicates distinct microbial community structures associated with these body sites. This clustering could reflect differences in environmental conditions, host-microbe interactions, or microbial functions between these sites[17].

The significant PERMANOVA test for all reported metrics (p-value 0.001) indicates significant differences between groups. The dispersion homogeneity test is also significant, indicating we cannot fully reject the null hypotesis (no differences between groups). The significance of the dispersion homogeneity test suggests that while there are clear differences between groups, the within-group variability must also be considered when interpreting these patterns.

Results of the Tukey test for multiple comparison of means is reported in Table 2. This provide deeper insights into the specific differences between microbial communities at different body sites.

**Table 2.**
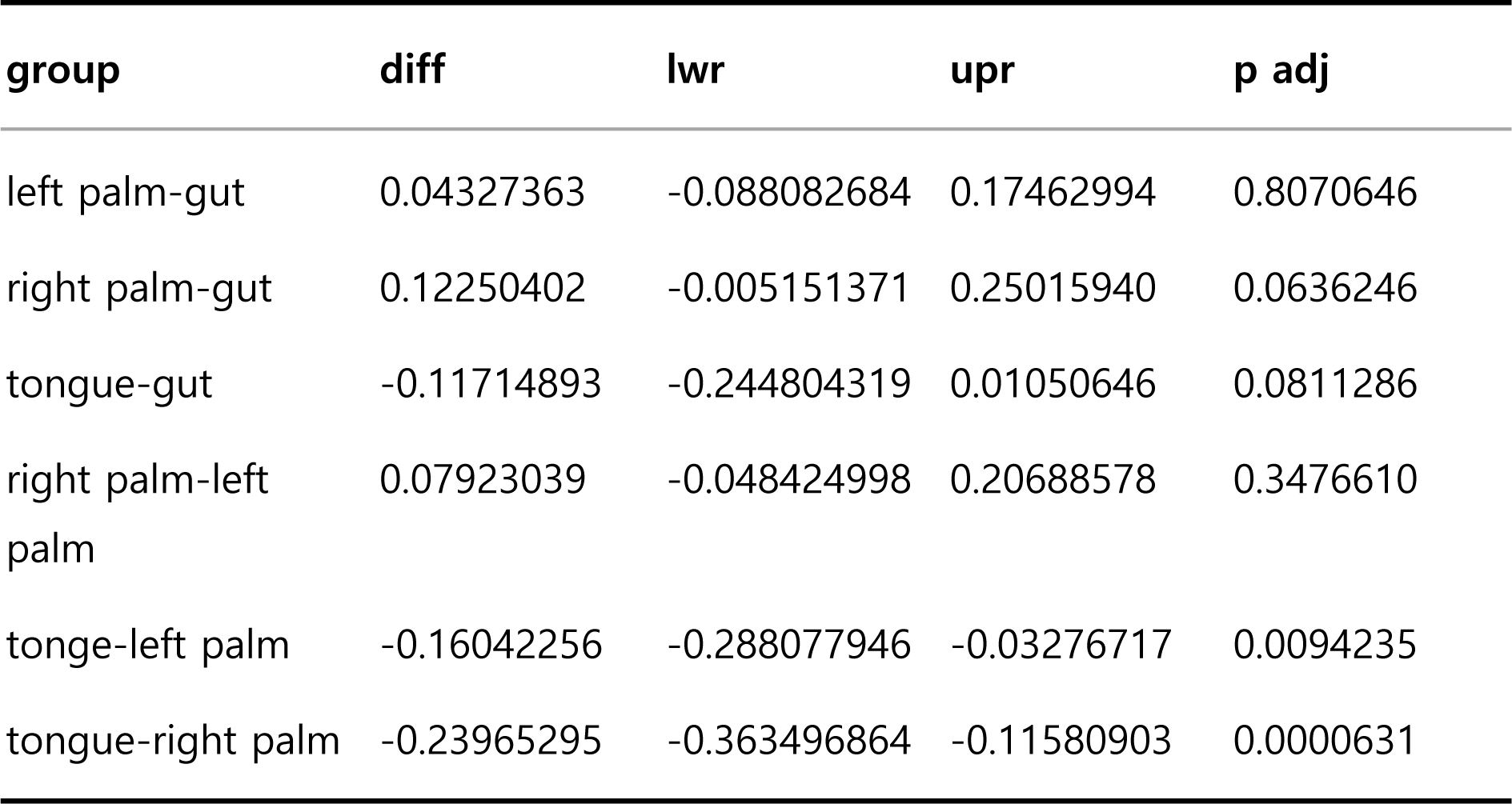
Tukey test for multiple comparisons of means.

| group | diff | lwr | upr | p adj |
| --- | --- | --- | --- | --- |
| left palm-gut | 0.04327363 | -0.088082684 | 0.17462994 | 0.8070646 |
| right palm-gut | 0.12250402 | -0.005151371 | 0.25015940 | 0.0636246 |
| tongue-gut | -0.11714893 | -0.244804319 | 0.01050646 | 0.0811286 |
| right palm-left palm | 0.07923039 | -0.048424998 | 0.20688578 | 0.3476610 |
| tonge-left palm | -0.16042256 | -0.288077946 | -0.03276717 | 0.0094235 |
| tongue-right palm | -0.23965295 | -0.363496864 | -0.11580903 | 0.0000631 |

### LEfSe Results and Biomarker Abundances

LEfSe analysis [13] identified key biomarkers differentially abundant across groups. *Bacteroidetes*, *Actinobacteria*, *Cyanobacteria*, *Proteobacteria*, and *Fusobacteria* were highlighted as significant biomarkers. Boxplots and heatmaps of biomarker abundances further illustrate the enrichment or depletion of these taxa in specific body sites, offering valuable insights into the functional roles of these microbial communities. These taxa are known for their varied metabolic capabilities and interactions with the host, suggesting their potential functional roles in maintaining health or contributing to disease states.

The Table 3 illustrates the results of the LEfSe analysis.

**Table 3.**
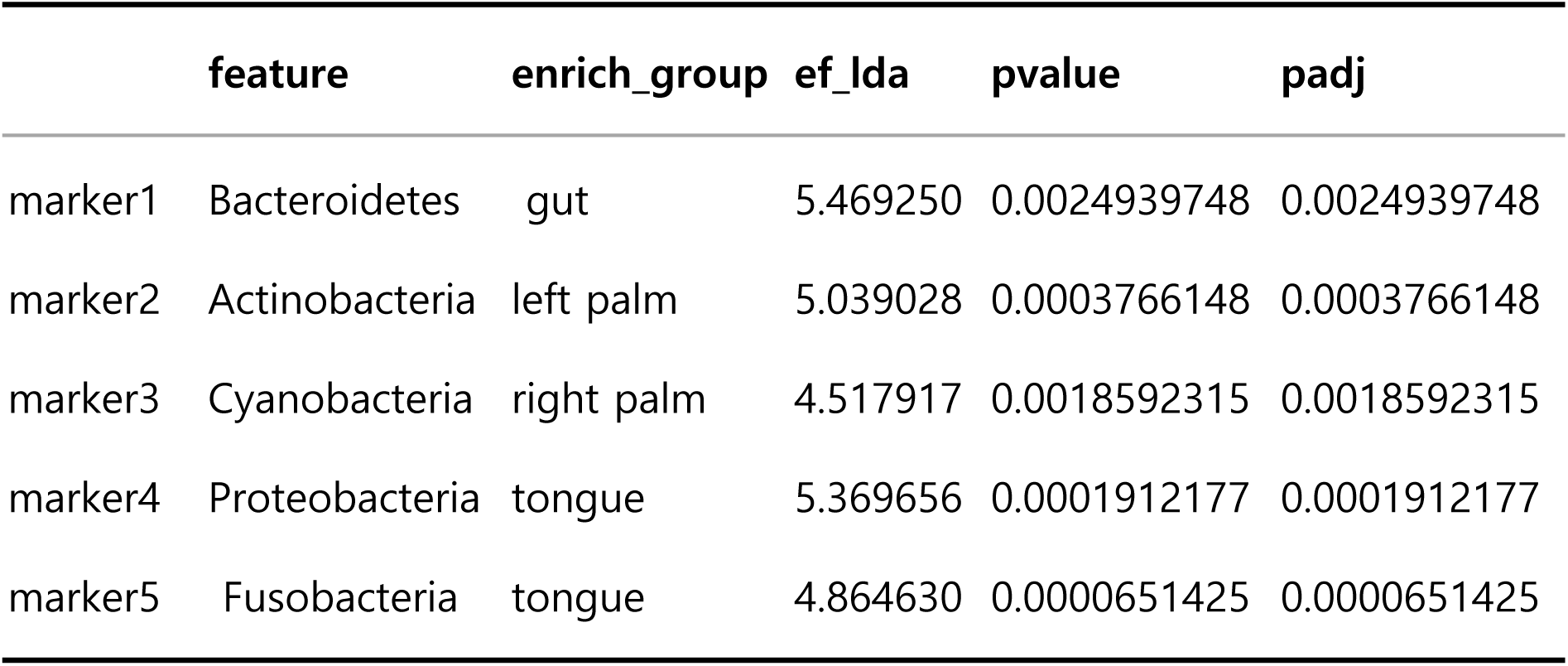
LEfSe analysis results.

|  | **feature** | **enrich_group** | **ef_lda** | **pvalue** | **padj** |
| --- | --- | --- | --- | --- | --- |
| marker1 | Bacteroidetes | gut | 5.469250 | 0.0024939748 | 0.0024939748 |
| marker2 | Actinobacteria | left palm | 5.039028 | 0.0003766148 | 0.0003766148 |
| marker3 | Cyanobacteria | right palm | 4.517917 | 0.0018592315 | 0.0018592315 |
| marker4 | Proteobacteria | tongue | 5.369656 | 0.0001912177 | 0.0001912177 |
| marker5 | Fusobacteria | tongue | 4.864630 | 0.0000651425 | 0.0000651425 |

The identification of key biomarkers differentially abundant across groups, as shown in Fig 16, provides insights into the functional roles of these microbial communities. The enrichment or depletion of specific taxa in certain body sites, as detailed in Figs 17 and 18, could have significant implications for understanding the microbiome’s contribution to health and disease.

**Fig 16.**
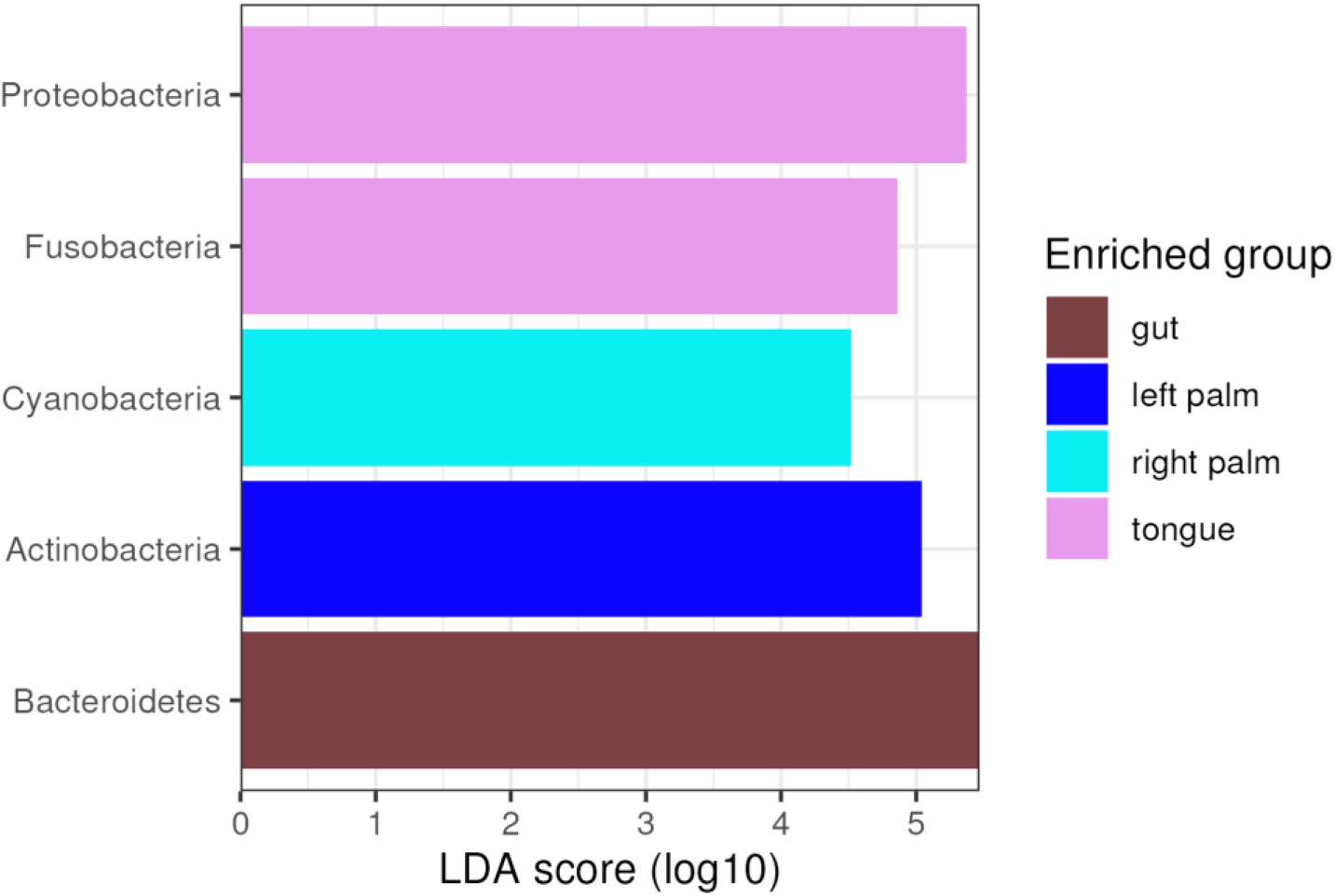
Differential Abundance Analysis with LEfSe: Barplot visualizing biomarkers identified through LEfSe analysis, indicating taxa that are significantly enriched or depleted in specific groups.

**Fig 17.**
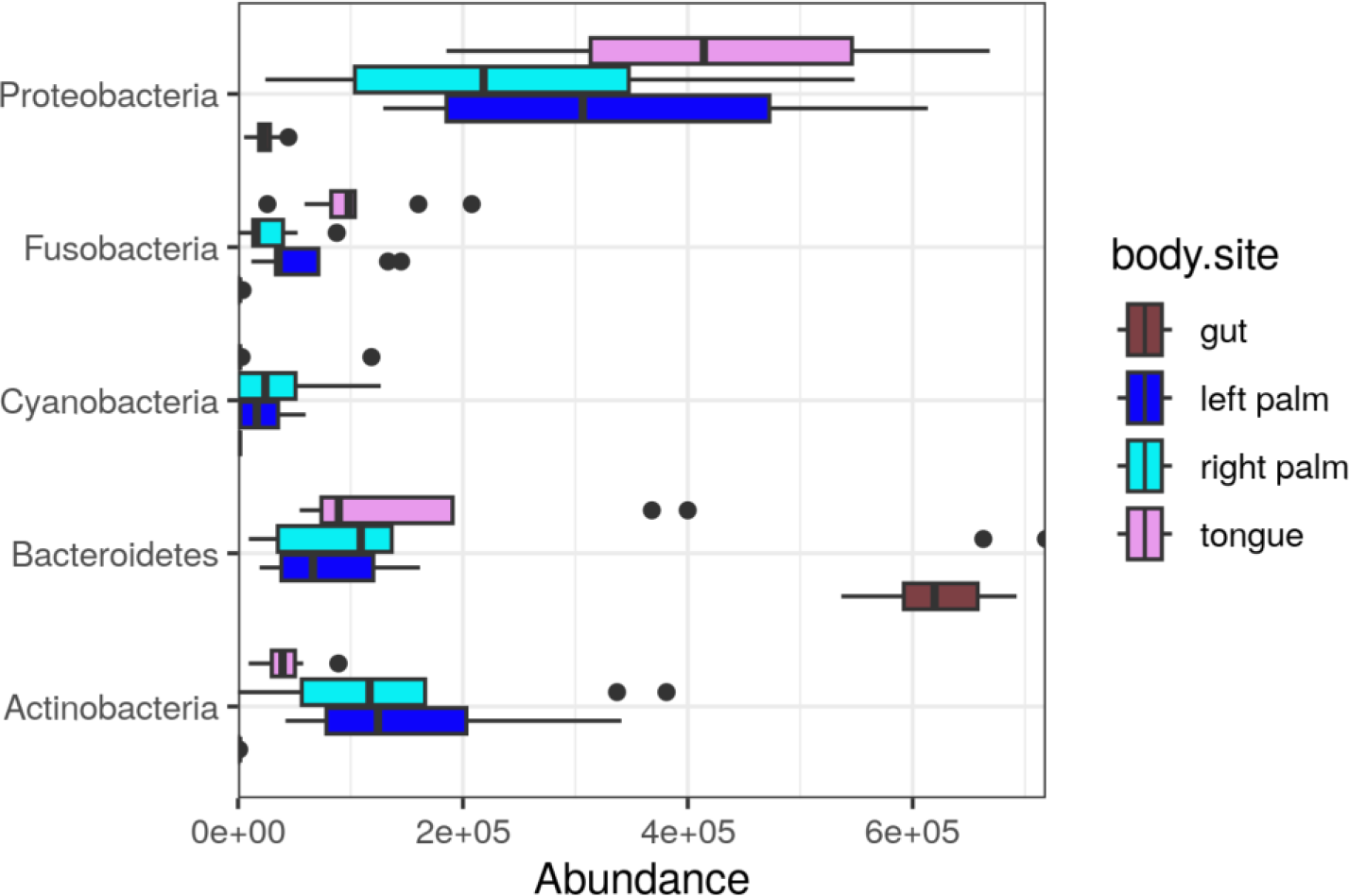
Visualization of Biomarker Abundance: Boxplots detailing the abundance of key biomarkers across groups, with gut samples enriched in *Bacteroidetes* and depleted in *Proteobacteria*, and vice versa for other groups.

**Fig 18.**
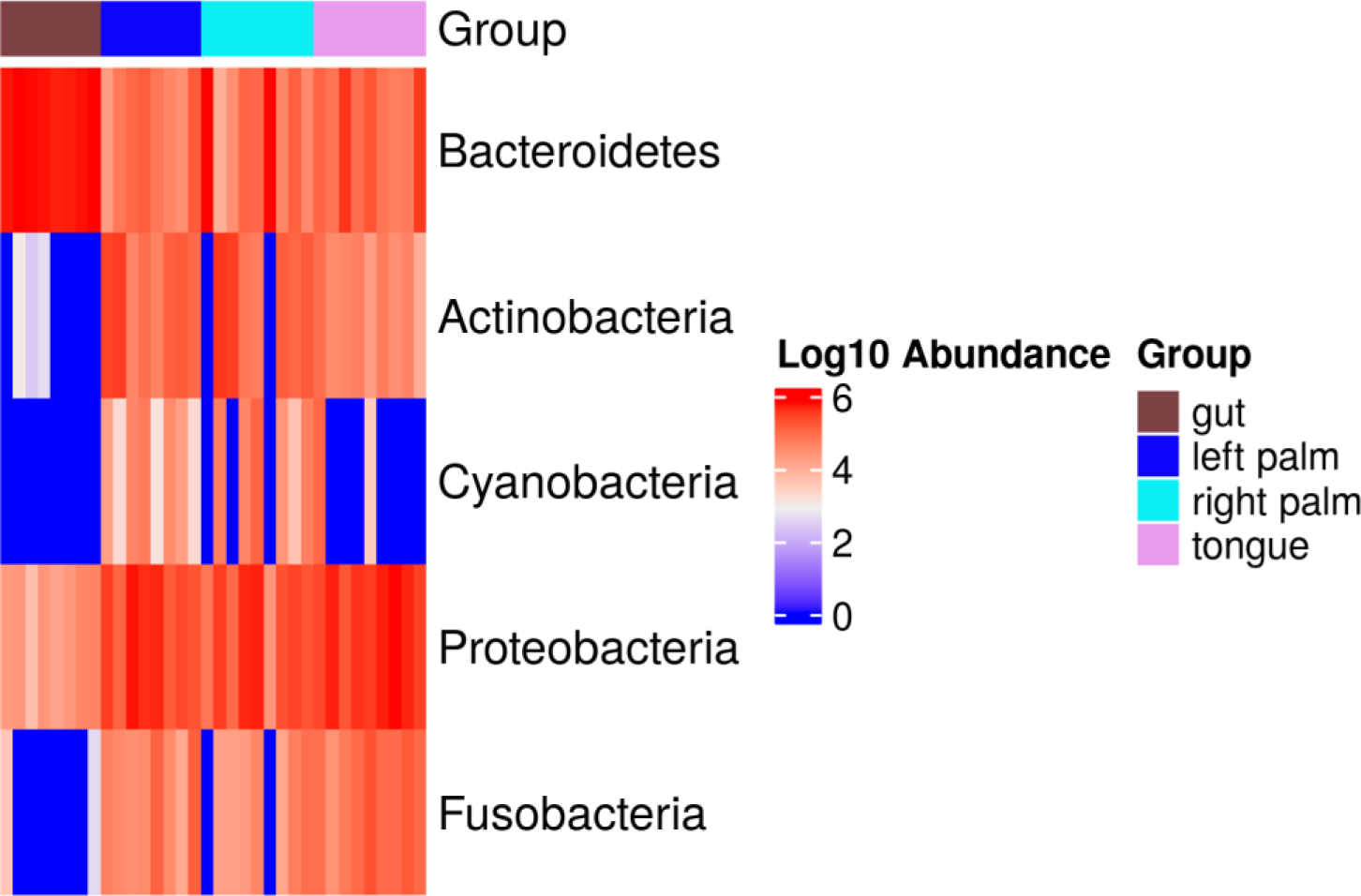
Visualization of Biomarker Abundance: Heatmap detailing the abundance of key biomarkers across groups, with gut samples enriched in *Bacteroidetes* and depleted in *Proteobacteria*, and vice versa for other groups.

For instance, the enrichment of Bacteroidetes in gut samples, as opposed to their depletion in samples where Proteobacteria are enriched, can indicate a shift in the microbial community structure that may influence gut health and disease. Bacteroidetes are known for their role in degrading complex molecules to produce energy, while Proteobacteria include a wide range of pathogens. The differential abundance of these taxa across body sites can thus offer insights into the microbial dynamics that underpin health and disease.

The functional roles of these microbial communities are of particular interest. For example, Actinobacteria are important for their antibiotic production, while Cyanobacteria play a role in nitrogen fixation. The presence or absence of these taxa in specific body sites can influence the local microbial ecosystem and, by extension, the host’s health. The enrichment of Fusobacteria in certain conditions, given their association with periodontal disease, could indicate a predisposition to or presence of disease states.

LEfSe analysis, as demonstrated in the provided figures and table, offers a powerful approach to uncovering the microbial taxa that define different body sites or conditions. By identifying and visualizing these key biomarkers, researchers can gain valuable insights into the composition and functional potential of microbial communities. This, in turn, can inform our understanding of the microbiome’s role in health and disease, guiding future research and potential therapeutic strategies.

## Conclusions

The exploration of microbiome data through the lens of MicrobiomePhylo has illuminated the intricate dynamics and diversity of microbial communities across various environments and host interactions. Key findings from utilizing this platform underscore the critical role of comprehensive, yet user-friendly, analytical tools in advancing our understanding of the microbiome [21]. By integrating robust statistical analyses with dynamic visualization capabilities, MicrobiomePhylo empowers researchers to uncover nuanced insights into microbial composition, diversity, and functional potential.

The potential impact of using MicrobiomePhylo for microbiome research is profound. For researchers, particularly those in the fields of bioinformatics and hospital research laboratories, the platform offers an unparalleled opportunity to delve deeper into the microbiome’s complexities. This can lead to groundbreaking discoveries in understanding disease mechanisms, developing novel therapeutic strategies, and enhancing personalized medicine approaches. Furthermore, the platform’s accessibility to users of varying computational expertise democratizes microbiome research, fostering a more inclusive scientific community [24].

Looking ahead, the continuous evolution of MicrobiomePhylo will be guided by the ever-expanding landscape of microbiome research. Future directions for the platform include the integration of metagenomic and metatranscriptomic data analysis capabilities [22], enhancing the breadth of microbial insights. Additionally, the development of more sophisticated machine learning models for predictive microbiome analysis could further revolutionize our approach to understanding and manipulating microbial communities for health and environmental benefits [23]

In conclusion, MicrobiomePhylo stands at the forefront of facilitating ground-breaking microbiome research. Its commitment to continuous improvement and adaptation to the latest scientific advancements ensures that it remains an invaluable resource for the research community. By empowering researchers with sophisticated tools for comprehensive microbiome analysis, MicrobiomePhylo is paving the way for novel discoveries and innovations that could have far-reaching implications for health, disease management, and environmental conservation..

## Limitations and Future Directions

While MicrobiomePhylo represents a significant advancement in microbiome data analysis, it is important to acknowledge the platform’s limitations and areas for future development. One limitation is the current focus on amplicon sequencing data, which, while powerful, provides only a snapshot of microbial community composition. Expanding the platform to include metagenomic, metatranscriptomic, and metabolomic data analysis would offer a more comprehensive view of microbial functions and interactions [25].

Another area for improvement is the enhancement of computational efficiency and scalability. As microbiome datasets grow in size and complexity, optimizing the platform’s algorithms and infrastructure to handle large-scale analyses efficiently will be crucial. This includes the implementation of cloud computing capabilities to facilitate the processing of extensive datasets without compromising performance [26].

Furthermore, the integration of longitudinal data analysis tools could significantly enhance the platform’s utility. The ability to track changes in microbial communities over time is essential for understanding the dynamics of hostmicrobe interactions and the impact of interventions on the microbiome [27].

Finally, fostering a collaborative community around MicrobiomePhylo could spur innovation and improvement. Encouraging user feedback, sharing analysis pipelines, and facilitating discussions on best practices can drive the platform’s evolution to meet the changing needs of the microbiome research community [24].

In conclusion, MicrobiomePhylo stands as a pivotal tool in the realm of microbiome research, offering powerful insights and fostering scientific discovery. By addressing current limitations and pursuing ambitious future directions, the platform is poised to remain at the forefront of this dynamic field, unlocking new horizons in our understanding of the microbial world.

## References

[1] Caporaso, J. G., Lauber, C. L., Walters, W. A., Berg-Lyons, D., Lozupone, C. A., Turnbaugh, P. J., … & Knight, R. (2011). Global patterns of 16S rRNA diversity at a depth of millions of sequences per sample. Proceedings of the National Academy of Sciences, 108(Supplement 1), 4516–4522.

[2] Bolyen, E., Rideout, J. R., Dillon, M. R., Bokulich, N. A., Abnet, C. C., Al-Ghalith, G. A., … & Caporaso, J. G. (2019). Reproducible, interactive, scalable and extensible microbiome data science using QIIME 2. Nature Biotechnology, 37(8), 852–857.

[3] McMurdie, P. J., & Holmes, S. (2013). phyloseq: an R package for reproducible interactive analysis and graphics of microbiome census data. PloS one, 8(4), e61217.

[4] Gilbert, J. A., Jansson, J. K., & Knight, R. (2014). The Earth Microbiome project: successes and aspirations. BMC biology, 12(1), 1–4.

[5] Anderson, M. J. (2001). A new method for non-parametric multivariate analysis of variance. Austral ecology, 26(1), 32–46.

[6] Lozupone, C., & Knight, R. (2005). UniFrac: a new phylogenetic method for comparing microbial communities. Applied and environmental microbiology, 71(12), 8228–8235.

[7] Schloss, P. D. (2024). Waste not, want not: Revisiting the analysis that called into question the practice of rarefaction. mSphere, 9(e00355-23). 10.1128/msphere.00355-23

[8] Weiss, S., Xu, Z. Z., Peddada, S., Amir, A., Bittinger, K., Gonzalez, A., … & Knight, R. (2017). Normalization and microbial differential abundance strategies depend upon data characteristics. Microbiome, 5(1), 27.

[9] Love, M. I., Huber, W., & Anders, S. (2014). Moderated estimation of fold change and dispersion for RNA-seq data with DESeq2. Genome biology, 15(12), 550.

[10] Benjamini, Y., & Hochberg, Y. (1995). Controlling the false discovery rate: a practical and powerful approach to multiple testing. Journal of the Royal Statistical Society. Series B (Methodological), 289–300.

[11] Chao, A., & Bunge, J. (2002). Estimating the number of species in a stochastic abundance model. Biometrics, 58(3), 531–539.

[12] Calle, M. L. (2019). Statistical Analysis of Metagenomics Data. Genomics & Informatics, 17(1), e6. doi: 10.5808/GI.2019.17.1.e6.

[13] Segata, N., Izard, J., Waldron, L., Gevers, D., Miropolsky, L., Garrett, W. S., & Huttenhower, C. (2011). Metagenomic biomarker discovery and explanation. Genome biology, 12(6), R60.

[14] Dunn, O. J. (1964). Multiple comparisons using rank sums. Technometrics, 6(3), 241–252.

[15] Wickham, H. (2016). ggplot2: Elegant Graphics for Data Analysis. Springer-Verlag New York.

[16] Human Microbiome Project Consortium. (2012). Structure, function and diversity of the healthy human microbiome. Nature, 486(7402), 207–214.

[17] Costello, E. K., Lauber, C. L., Hamady, M., Fierer, N., Gordon, J. I., & Knight, R. (2009). Bacterial community variation in human body habitats across space and time. Science, 326(5960), 1694–1697.

[18] Kuczynski, J., Lauber, C. L., Walters, W. A., Parfrey, L. W., Clemente, J. C., Gevers, D., & Knight, R. (2012). Experimental and analytical tools for studying the human microbiome. Nature Reviews Genetics, 13(1), 47–58.

[19] Caporaso, J. G., Lauber, C. L., Walters, W. A., Berg-Lyons, D., Huntley, J., Fierer, N., Owens, S. M., Betley, J., Fraser, L., Bauer, M., Gormley, N., Gilbert, J. A., Smith, G., & Knight, R. (2012). Ultra-high-throughput microbial community analysis on the Illumina HiSeq and MiSeq platforms. The ISME Journal, 6(8), 1621–1624.

[20] Legendre, P., & Legendre, L. (2012). Numerical ecology. Elsevier.

[21] Gilbert, J. A., Blaser, M. J., Caporaso, J. G., Jansson, J. K., Lynch, S. V., & Knight, R. (2018). Current understanding of the human microbiome. Nature Medicine, 24(4), 392–400.

[22] Quince, C., Walker, A. W., Simpson, J. T., Loman, N. J., & Segata, N. (2017). Shotgun metagenomics, from sampling to analysis. Nature Biotechnology, 35(9), 833–844.

[23] Knights, D., Costello, E. K., & Knight, R. (2011). Supervised classification of human microbiota. FEMS Microbiology Reviews, 35(2), 343–359.

[24] Scholz, M., Ward, D. V., Pasolli, E., Tolio, T., Zolfo, M., Asnicar, F., … & Segata, N. (2016). Strain-level microbial epidemiology and population genomics from shotgun metagenomics. Nature Methods, 13(5), 435–438.

[25] Knight, R., Vrbanac, A., Taylor, B.C., Aksenov, A., Callewaert, C., Debelius, J., … & McDonald, D. (2018). Best practices for analysing microbiomes. Nature Reviews Microbiology, 16(7), 410–422.

[26] Sczyrba, A., Hofmann, P., Belmann, P., Koslicki, D., Janssen, S., Dröge, J., … & McHardy, A.C. (2017). Critical Assessment of Metagenome Interpretation—a benchmark of metagenomics software. Nature Methods, 14(11), 1063–1071.

[27] Faust, K., Lahti, L., Gonze, D., de Vos, W.M., & Raes, J. (2015). Metagenomics meets time series analysis: unraveling microbial community dynamics. Current Opinion in Microbiology, 25, 56–66.

[28] Grice, E.A., & Segre, J.A. “The skin microbiome.” Nature Reviews Microbiology 9.4 (2011): 244–253.

[29] Ley, R.E., et al. “Microbial ecology: human gut microbes associated with obesity.” Nature 444.7122 (2006): 1022–1023.

[30] Findley, K., & Grice, E.A. “The skin microbiome: A focus on pathogens and their association with skin disease.” PLoS Pathogens 10.11 (2014): e1004436.

[31] Flint, H.J., Scott, K.P., Louis, P., & Duncan, S.H. “The role of the gut microbiota in nutrition and health.” Nature Reviews Gastroenterology & Hepatology 9.10 (2012): 577–589.

[32] Frank, D.N., et al. “Molecular-phylogenetic characterization of microbial community imbalances in human inflammatory bowel diseases.” Proceedings of the National Academy of Sciences 104.34 (2007): 13780–13785.

[33] Turnbaugh, P.J., et al. “An obesity-associated gut microbiome with increased capacity for energy harvest.” Nature 444.7122 (2006): 1027–1031.

[34] Belkaid, Y., & Hand, T.W. “Role of the microbiota in immunity and inflammation.” Cell 157.1 (2014): 121–141.

[35] Haiser, H.J., & Turnbaugh, P.J. “Is it time for a metagenomic basis of therapeutics?” Science 336.6086 (2012): 1253–1255.

[36] Turnbaugh, P.J., et al. “The human microbiome project.” Nature 449, 804–810 (2007).

[37] Costello, E.K., et al. “Bacterial community variation in human body habitats across space and time.” Science 326, 1694–1697 (2009).

[38] Lozupone, C.A., et al. “Diversity, stability and resilience of the human gut microbiota.” Nature 489, 220–230 (2012).

[39] Hill, C., et al. “The International Scientific Association for Probiotics and Prebiotics consensus statement on the scope and appropriate use of the term probiotic.” Nature Reviews Gastroenterology & Hepatology 11, 506–514 (2014).

[40] Kumar, H., et al. (2012). “Diversity, functionality and health benefits of Lactobacilli and Streptococci.” Trends in Food Science & Technology, 27(2), 98–107.

[41] Wexler, H. M. (2007). “Bacteroides: the good, the bad, and the nitty-gritty.” Clinical Microbiology Reviews, 20(4), 593–621.

[42] Post, D. M. B., et al. (2019). “The Complex Role of Haemophilus influenzae in Chronic Obstructive Pulmonary Disease.” Frontiers in Immunology, 10, 2340.

[43] Grice, E. A., & Segre, J. A. (2011). “The skin microbiome.” Nature Reviews Microbiology, 9(4), 244–253.

[44] Shin, N. R., Whon, T. W., & Bae, J. W. (2015). “Proteobacteria: microbial signature of dysbiosis in gut microbiota.” Trends in Biotechnology, 33(9), 496–503.

[45] Gloor, G. B., Macklaim, J. M., Pawlowsky-Glahn, V., & Egozcue, J. J. (2017). Microbiome Datasets Are Compositional: And This Is Not Optional. Frontiers in Microbiology, 8, 2224.

[46] Aitchison, J. (1986). The Statistical Analysis of Compositional Data. Chapman & Hall Ltd.

[47] Wilkinson, L., & Friendly, M. (2009). The history of the cluster heat map. The American Statistician, 63(2), 179–184.

[48] Qin, J., Li, R., Raes, J., Arumugam, M., Burgdorf, K. S., Manichanh, C., … & Wang, J. (2010). A human gut microbial gene catalogue established by meta-genomic sequencing. Nature, 464(7285), 59–65.

[49] Shade, A., Peter, H., Allison, S. D., Baho, D. L., Berga, M., Bürgmann, H., … & Martiny, J. B. H. (2012). Fundamentals of microbial community resistance and resilience. Frontiers in Microbiology, 3, 417.

[50] Heintz-Buschart, A., & Wilmes, P. (2018). Human gut microbiome: function matters. Trends in Microbiology, 26(7), 563–574.

[51] Le Chatelier, E., Nielsen, T., Qin, J., Prifti, E., Hildebrand, F., Falony, G., … & Ehrlich, S. D. (2013). Richness of human gut microbiome correlates with metabolic markers. Nature, 500(7464), 541–546.

